# Inhibitory basal ganglia nuclei differentially innervate pedunculopontine nucleus subpopulations and evoke differential motor and valence behaviors

**DOI:** 10.1101/2024.08.05.606694

**Authors:** Michel Fallah, Kenea C Udobi, Aleksandra E Swiatek, Chelsea B Scott, Rebekah C Evans

**Affiliations:** Interdisciplinary Program in Neuroscience, Georgetown University, Washington, DC, USA 20007; Department of Neuroscience, Georgetown University Medical Center, Washington, DC, USA 20007

**Keywords:** basal ganglia, brainstem, locomotion, reward, mesencephalic locomotor region, inhibition, electrophysiology, optogenetics, substantia nigra, pedunculopontine nucleus

## Abstract

The canonical basal ganglia model predicts that the substantia nigra *pars reticulata* (SNr) and the globus pallidus *externa* (GPe) will have specific effects on locomotion: the SNr inhibiting locomotion and the GPe enhancing it. In this study, we use *in vivo* optogenetics to show that the GPe exerts non-canonical effects on locomotion while the SNr has no gross motor impact through inhibition of the PPN. We also show that these structures mediate opposing effects on reward. To understand how these structures differentially modulate the PPN, we use *ex vivo* whole-cell recording with optogenetics to comprehensively dissect the SNr and GPe synaptic connections to regionally- and molecularly-defined populations of PPN neurons. The SNr inhibits all PPN subtypes, but most strongly inhibits caudal glutamatergic neurons. The GPe selectively inhibits caudal glutamatergic and GABAergic neurons, avoiding both cholinergic and rostral cells. This circuit characterization reveals non-canonical basal ganglia pathways for locomotion and valence.

**Graphical Abstract:** 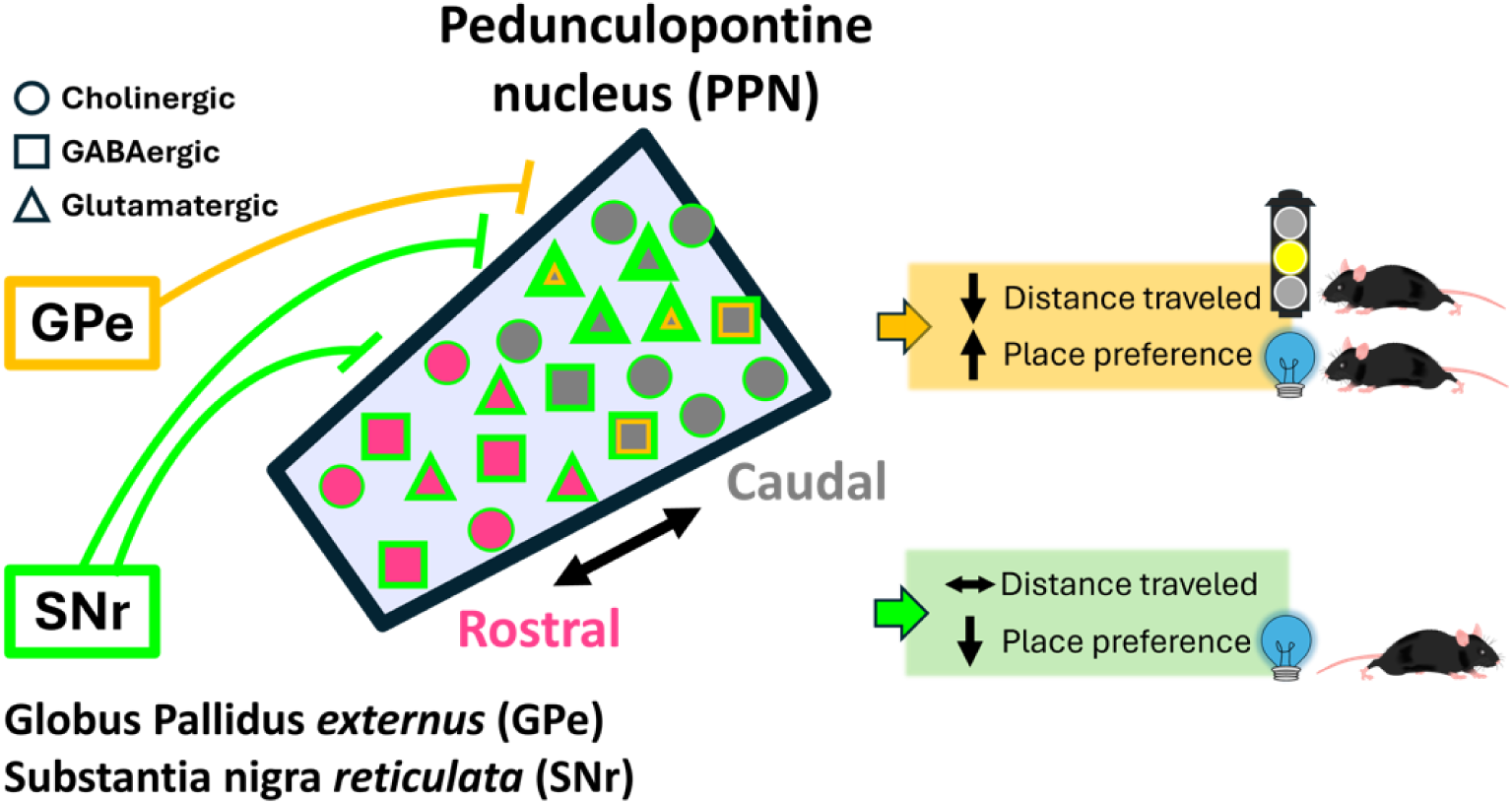

## Introduction

The pedunculopontine nucleus (PPN) is a brainstem structure heavily interconnected with subcortical structures, such as the basal ganglia, thalamus, and spinal cord. The PPN has been implicated in locomotor control and valence processing^1–4^. However, understanding PPN circuitry is complicated by its anatomical and molecular heterogeneity. It has three molecularly-defined cell types, distinguished by their major neurotransmitter^5^ and distinct rostral and caudal topography. While cholinergic and glutamatergic neurons are more densely packed in the caudal PPN, GABAergic neurons are more densely packed in the rostral PPN^5,6^. These molecularly-defined populations display very minimal overlap^5,7,8^. This unique topography has been conserved across species^9–11^, but it is unknown how the basal ganglia connects to each of the distinct cell types in the rostral and caudal PPN regions to modulate locomotor and valence outputs.

While the PPN interacts with all basal ganglia nuclei^3,9^, the inhibitory inputs from the substantia nigra *pars reticulata* (SNr) and the globus pallidus *externus* (GPe) to the PPN are of particular interest because of their roles in the canonical motor output pathways for ‘stop’ and ‘go,’ respectively. Both the SNr and GPe are known to send axonal projections to the PPN^12–18^. Previous studies have identified SNr and GPe monosynaptic connections to the PPN cholinergic and glutamatergic neurons using rabies tracing^13–17^. However, inputs to the GABAergic neurons have only been described in terms of the entire mesencephalic locomotor region (MLR), which extends beyond the PPN^14^. While SNr projections to MLR GABAergic neurons were identified but cannot be isolated to the PPN, GPe projections to the MLR GABAergic neurons were not detected^14^. Of note, these previous studies consider the PPN as a whole rather than separating its rostral and caudal subregions. While viral tracing can provide insight into neuroanatomical connections, limitations of these techniques^19–21^ and evidence that the rostral and caudal PPN may have distinct functions^22–25^ prompt a need for electrophysiological studies to determine whether the SNr and GPe inputs differentially innervate regionally-defined PPN neurons.

Optogenetic manipulations of the molecularly-defined PPN cell types have resulted in variable, and sometimes contradictory, locomotor and valence behaviors^7,14,16,17,22–33^. More recent studies have refined our understanding of these opposing behavioral effects by selectively stimulating the rostral or caudal GABAergic and glutamatergic PPN neurons^22–25^. In the basal ganglia, direct stimulation of molecularly-defined SNr and GPe subpopulations can also have distinct effects on locomotion and valence^34–42^. Despite the potential for the basal ganglia to modify behavior through the motor brainstem^14^, it is not clear how the SNr and GPe influence the rostral and caudal PPN to modify locomotion and valence.

Here, we use *ex vivo* whole-cell patch clamp electrophysiology paired with optogenetics to selectively stimulate the SNr or GPe axons while recording inhibitory inputs to the cholinergic, GABAergic, and glutamatergic PPN neurons to comprehensively characterize the strength and pattern of inhibition across the rostrocaudal PPN axis. We find that the GPe preferentially inhibits caudal GABAergic and glutamatergic PPN neurons whereas the SNr most strongly inhibits a caudal ‘hotspot’ of PPN glutamatergic neurons. Stimulating SNr or GPe axons over the PPN *in vivo* evokes opposing valence processing outcomes with place aversion during SNr stimulation and place preference during GPe stimulation. Surprisingly, and counter to the predictions of the canonical basal ganglia model, the GPe decreases locomotion through its projections to the PPN. While both the SNr and GPe inhibit the PPN, our results show that each nucleus differentially modulates the activity of specific cell types in the rostral and caudal PPN and are implicated in non-canonical basal ganglia circuits for modulating locomotion and valence processing.

## Results

### SNr and GPe axons display distinct distribution patterns across the rostral and caudal PPN

To characterize the axon distribution pattern from the SNr and GPe throughout the rostral and caudal extent of the PPN, we stereotaxically injected a channelrhodopsin virus with a fluorescent marker (AAV1-hSyn-ChR2-eYFP) into either the SNr or GPe of ChAT-Cre/Ai9-tdTomato mice (**Figure 1A&C**). We sliced 200 μm sagittal slices from mouse brains three weeks after viral injection, and cleared the brain slices using the CUBIC clearing method^43^. Using the red fluorescent ChAT-positive neurons to define the borders of the PPN, we found that SNr axons appear in both the rostral and caudal PPN. The SNr axons fill the rostral PPN evenly, but appear more distinctly clustered in specific areas of the caudal PPN (**Figure 1B**), whereas GPe axonal projections more densely fill the caudal region and strikingly avoid the rostral region (**Figure 1D**). These anatomical findings predict that the SNr will inhibit both rostral and caudal PPN neurons, while the GPe will selectively inhibit the caudal PPN. However, this axon imaging method does not reveal whether these axons form functional synapses with the neurons in the PPN and does not provide information on the strength of inhibition from each structure onto specific PPN cell types. Therefore, to evaluate the functional connectivity and synaptic characteristics from these inhibitory basal ganglia output structures to regionally- and molecularly-defined PPN neural subpopulations, we used *ex vivo* whole-cell electrophysiology to systematically record the inhibitory input to the rostral and caudal PPN from the SNr and GPe in three genetic mouse lines (ChAT-cre, Vgat-cre, and Vglut2-cre) which label cholinergic, GABAergic, and glutamatergic PPN subpopulations, respectively (**Figure S1**).

**Figure 1.**
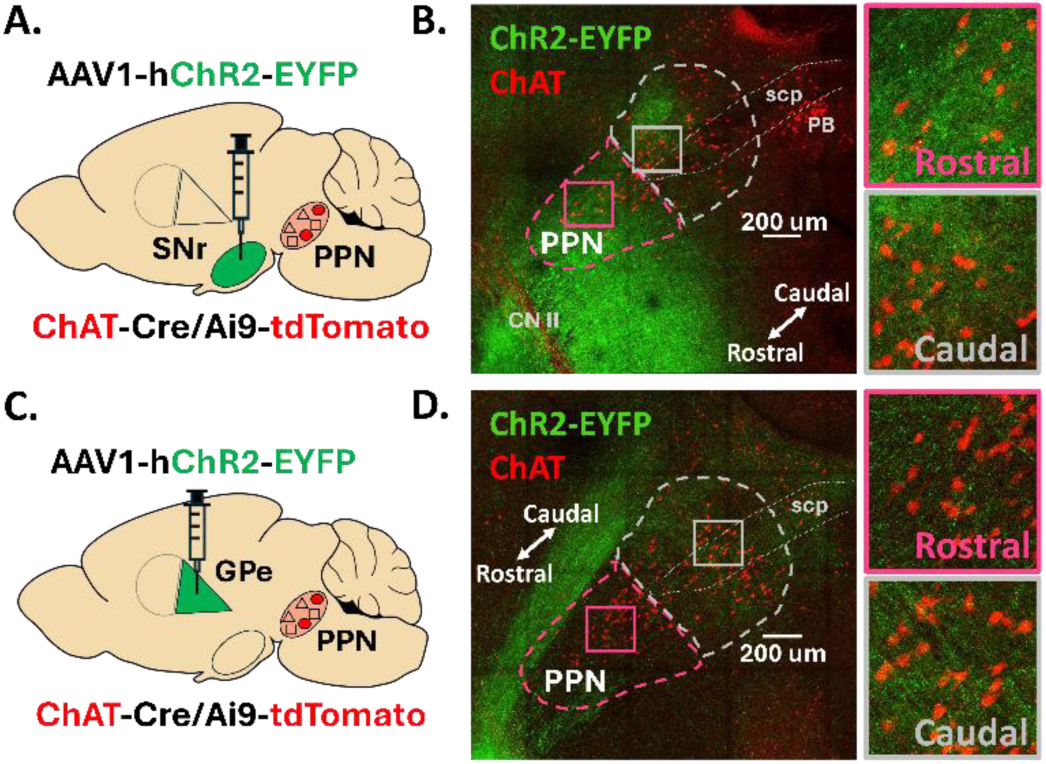
SNr and GPe axons display distinct distribution patterns across the rostral and caudal PPN. (A,C) Stereotaxic injection of AAV1 delivering *hSyn-ChR2-eYFP* to the SNr or GPe of ChAT-Cre/Ai9-tdTomato mice, respectively. (B, D) Confocal images of EYFP-filled SNr or GPe axons across the PPN, respectively. *CNII: cranial nerve II, scp: superior cerebellar peduncle; PB: parabrachial nucleus*

### The SNr inhibits cholinergic neurons in both the rostral and caudal PPN to a similar extent, but evokes stronger rebound in rostral neurons

To evaluate the functional connection between the SNr and cholinergic PPN neurons, we performed *ex vivo* whole-cell patch clamp electrophysiology paired with optogenetics to record synaptic currents in ChAT-Cre/Ai9-tdTomato mice injected with ChR2 in the SNr three weeks prior (**Figure 2A**). Neurons were targeted for whole-cell patch clamp in both the rostral and caudal PPN across three sagittal slices to evaluate the medial to lateral extent of the PPN (**Figure 2B**). To measure optically-evoked inhibitory post-synaptic currents (oIPSCs), cholinergic PPN neurons were held at −50 mV in whole-cell voltage clamp and blue (470 nm) light was applied (2 ms pulse duration, 13 mW) to activate the ChR2-infected SNr axons. In connected neurons, inhibitory currents were recorded in the presence of glutamatergic receptor blockers (AP5, NBQX, CNQX) during stimulation (**Figure 2C**). In a subset of neurons, a GABA-a receptor blocker, GABAzine, was applied to confirm the oIPSCs were GABA-mediated (**Figure 2C**). GABAzine eliminated SNr-evoked inhibitory currents in most cells, but a small residual current remained in some cells (see *Methods* for details). We found that all rostral (n=19/19 cells, N=6 mice) and essentially all caudal (n=26/27, N=6) recorded cholinergic PPN neurons responded to SNr axon activation with an observable oIPSC (**Figure 2D**).

**Figure 2.**
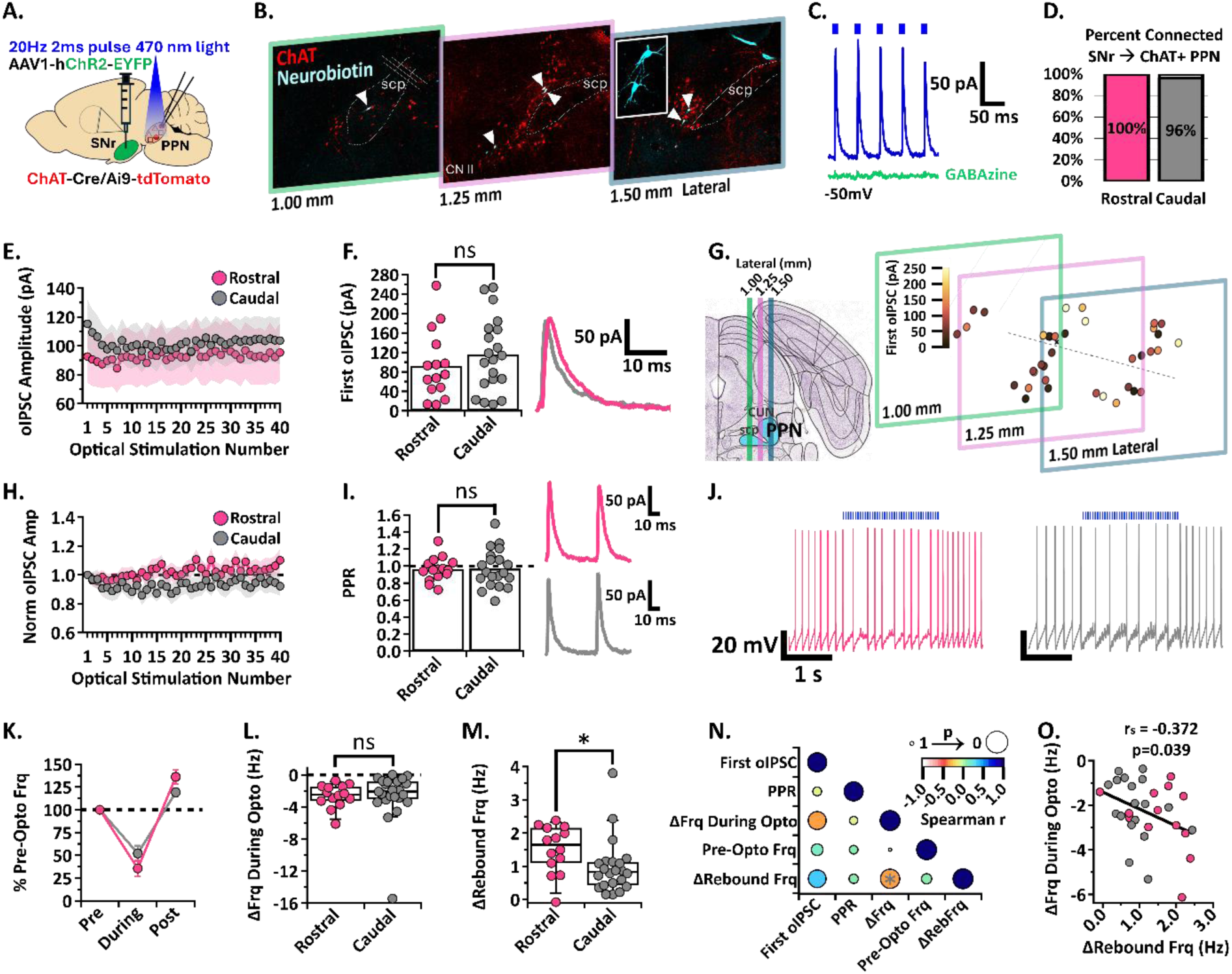
SNr inhibition of rostral and caudal ChAT+ PPN neurons. (A) Experimental set up to identify red ChAT+ PPN neurons for whole-cell patch clamp while stimulating ChR2-filled SNr axons [N=6]. (B) White arrowheads pointing to neurobiotin-filled patched neurons within the PPN across three 200µm slices. (C) Example trace of the first five oIPSCs [blue] in the 2-second 20 Hz train inhibited by GABA-a receptor blocker, GABAzine [green], while holding the cell at −50mV. (D) Percent connected among patched neurons in the rostral and caudal regions. (E) Average oIPSC amplitude at each of 40 optogenetic light pulses in n=15 rostral neurons and n=20 caudal neurons. (F) *Left*, Individual cell data for the first oIPSC amplitude and, *right*, example current traces. (G) Cell mapping of patched neuron locations with the first oIPSC amplitude represented by the color scale. (H) Normalized current amplitudes in E. (I) *Left*, Individual cell data for the PPR between the first two oIPSC amplitudes in the train and, *right*, example current traces. (J) Example voltage traces of action potential firing during a 2-second 20 Hz train stimulation in rostral (left) and caudal (right) neurons. (K) Percent of pre-optical stimulation firing frequency during stimulation and rebound in n=14 rostral vs n=23 caudal neurons. (L) Individual cell data for the absolute change in frequency during optical stimulation [ΔFrq During Opto]. (M) Individual cell data for the absolute change in rebound frequency post-stimulation [ΔRebFrq]; rostral vs. caudal p=0.0142. (N) Correlation analysis, color scale representing Spearman r [-1,1] and size representing p-value [1,0]. (O) Negative correlation between the absolute change in frequency during stimulation and post-stimulation rebound; r=-0.372, p=0.039. * p<0.05; bar graph data represent mean ± SEM; box plots show median line with boxes showing IQR and whiskers showing 9th and 91st percentiles.

To determine whether the SNr input to the rostral and caudal PPN cholinergic neurons differed in synaptic strength, we measured the amplitude of each oIPSC in a 20 Hz train of 2 ms optical stimulations applied for 2 seconds. There was no difference in amplitude of the first oIPSC in the stimulus train between rostral vs caudal cholinergic neurons (mean ± SEM; first oIPSC n=15 Rostral: 92.6 ± 18 pA, n=20 Caudal: 115.2 ± 17 pA; unpaired t-test, t=0.9041, df=33, p=0.3725; **Figure 2E&F**). Going forward, the first oIPSC amplitude of the train will be referred to as the first oIPSC.

While the rostral and caudal anatomical separation of PPN neurons is commonly used, it is a relatively coarse division. Therefore, we carefully mapped the location of each recorded cholinergic PPN neuron by matching slices to their bregma reference in the Paxinos and Franklin’s mouse brain atlas and aligning the midline of cholinergic neuron distribution which separates the loosely spread rostral neurons and densely packed caudal neurons (See *Methods* for details, **Figure 2G**). The strength of the SNr inhibitory connection is depicted by a color scale representative of the first oIPSC (**Figure 2G, *right***). Throughout both the rostral and caudal PPN, cholinergic neurons receiving larger inputs from the SNr are intermixed with neurons receiving smaller inputs.

To compare short-term synaptic plasticity characteristics between rostral and caudal cholinergic PPN neurons, we normalized the amplitude of each current in the train to the first current. The oIPSC amplitude remained relatively constant with each subsequent pulse in both rostral and caudal cholinergic neurons (**Figure 2H**). In addition, the paired-pulse ratio (PPR) of the peak of the first two currents in the stimulation train did not differ (mean ± SEM; PPR n=15 Rostral: 0.96 ± 0.04, n=20 Caudal: 0.97 ± 0.05; unpaired t-test, t=0.1082, df=33, p=0.9145; **Figure 2I**). SNr synaptic inputs onto cholinergic neurons do not undergo significant short-term synaptic plasticity in either the rostral or caudal PPN.

Although oIPSC amplitude is indicative of the strength of a synapse, current size does not always correlate with functional impact on a neuron’s action potential output^20,44,45^. Because cholinergic PPN neurons display spontaneous firing^46–50^, we characterized the impact of SNr-mediated inhibition on action potential firing rate. While optically stimulating SNr axons with the same 20 Hz blue light protocol as above, we observed a decrease in firing rate in both rostral and caudal cholinergic PPN neurons **Figure 2J-K**). However, there was no significant difference between the percent frequency change in rostral and caudal neurons during SNr axon stimulation (mean ± SEM; %Frequency n=14 Rostral: During Stimulation 35.62 ± 8.87 %, Post Stimulation 136.1 ± 7.803 %; n=23 Caudal: During Stimulation 52.11 ± 8.43 %, Post Stimulation 119.1 ± 4.614 %; 2way ANOVA, Stimulation_F(1, 35)=81.9_ p<0.0001, Region_F(1, 35)=0.002052_ p=0.9641, Interaction_F(1, 35)=3.273_ p=0.0790; **Figure 2K**) or in the absolute decrease in firing frequency during optical stimulation [median (IQR); ΔFrq During Opto n=14 Rostral: −2.47 Hz (−3.37 to −1.44 Hz), n=23 Caudal: −2.10 Hz (−3.11 to −0.84 Hz); Mann Whitney, U=131, p=0.3602; **Figure 2L**]. These data show that the SNr equally inhibits action potential firing in rostral and caudal PPN cholinergic neurons.

Post-inhibitory rebound firing has been observed in the PPN as well as other brainstem structures^49,51–57^ and is associated with greater excitability, temporal encoding, and generation of oscillatory activity^58^. Indeed, we found that the rostral and caudal cholinergic PPN neurons exhibited an increase in action potential firing after inhibition (**Figure 2K**). This is in agreement with previous literature showing rebound spikes following inhibitory postsynaptic potentials recorded in PPN neurons of male rats during electrical stimulation of the SNr^51,52^. However, rostral cholinergic neurons had greater rebound activity compared with caudal cholinergic neurons as measured by the absolute change in rebound frequency [median (IQR); ΔRebFrq n=14 Rostral: 1.64 Hz (1.00 to 2.17 Hz), n=22 Caudal: 0.83 Hz (0.42 to 1.14 Hz); Mann Whitney, U=79, p=0.0142; **Figure 2M**]. This aligns with previous findings showing that rostral cholinergic neurons tend to have stronger intrinsic rebound activity compared to caudal neurons^49^. Importantly, we show that such rebound can be evoked by synaptic inhibition from the SNr.

To detect trends among SNr connectivity characteristics and intrinsic electrophysiological properties of cholinergic PPN neurons, we performed a correlation analysis to determine the strength and directionality of the relationship among five parameters [first oIPSC, PPR, ΔFrq During Opto, pre-optical stimulation frequency (Pre-Opto Frq), and ΔRebFrq; **Figure 2N**]. We found a weak but significant negative correlation between the frequency decrease during SNr axon stimulation and the rebound frequency increase (Spearman r=-0.372, p=0.039, **Figure 2N&O**). This finding is expected since strong inhibition is more effective at producing rebound activity than weak inhibition. However, rostral PPN cholinergic neurons do not receive significantly stronger inhibition from the SNr compared with caudal cholinergic neurons (**Figure 2F&L**). Therefore, larger rebound frequency in rostral neurons is likely due to differences in intrinsic cellular properties rather than synaptic properties^49^. Put together, these findings show that the SNr inhibits rostral and caudal cholinergic PPN neurons to a similar extent, but evokes stronger rebound firing in the rostral neurons.

### The SNr inhibits GABAergic neurons in both the rostral and caudal PPN to a similar extent

While there are several rabies tracing studies suggesting SNr axons project to the cholinergic and glutamatergic PPN neurons^13–17^, SNr projections to the GABAergic PPN neurons are inferred from a study of the entire mesencephalic locomotor region^14^ and a characterization of their synaptic strength has not been conducted. Therefore, we evaluated the synaptic characteristics between the SNr and GABAergic neurons in the rostral and caudal PPN. In Vgat-Cre/Ai9-tdTomato mice, we injected ChR2 into the SNr to optically stimulate axons over the PPN while patching Vgat+ GABAergic PPN neurons (**Figure 3A**). We post-hoc stained the slices for ChAT+ neurons to verify the location of all patched GABAergic neurons within the cholinergic PPN. We found that all rostral (n=12/12, N=6) and most caudal (n=20/22, N=6) recorded GABAergic PPN neurons received inhibitory input from the SNr (**Figure 3B**).

**Figure 3.**
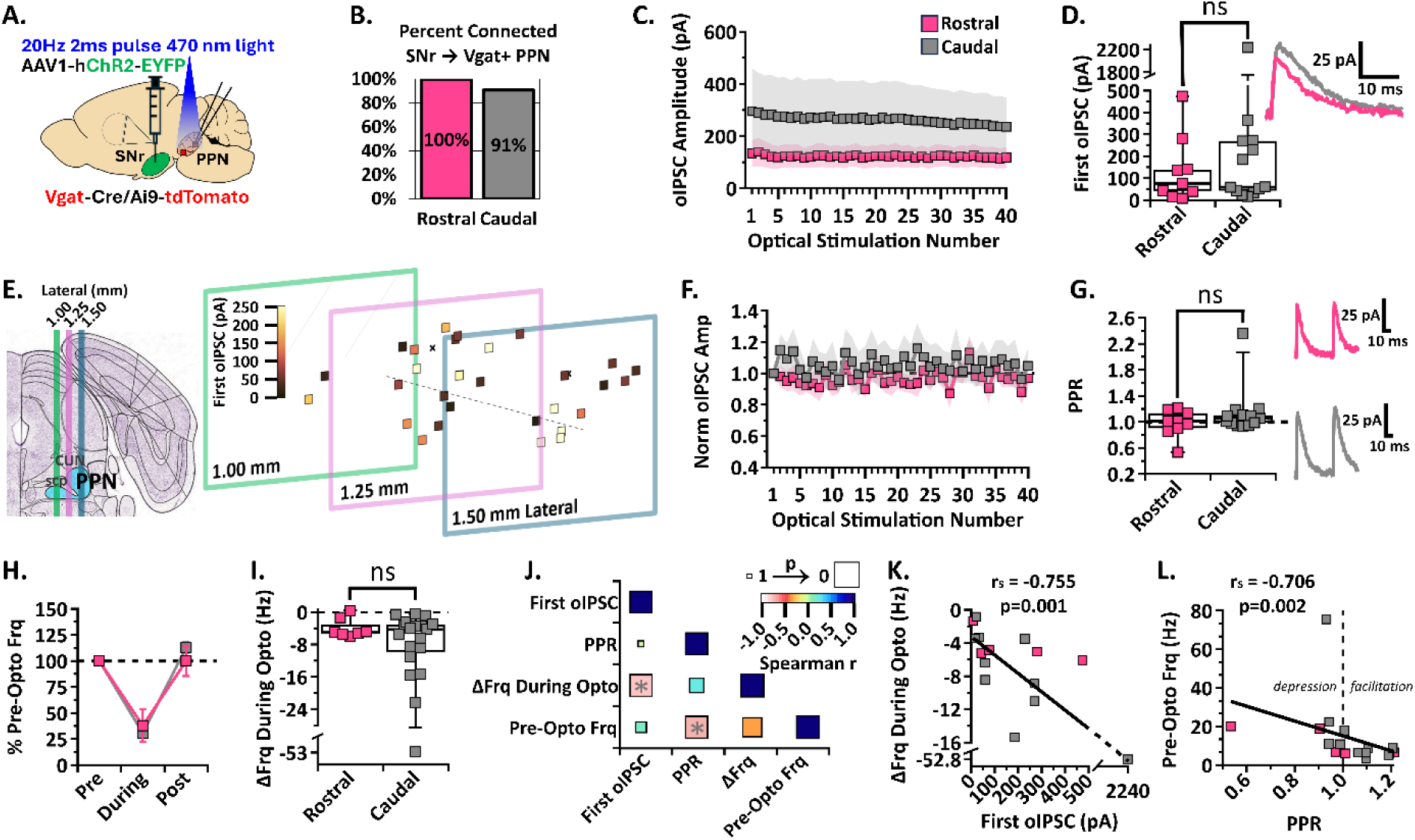
SNr inhibition of rostral and caudal Vgat+ PPN neurons. (A) Experimental set up to identify red Vgat+ PPN neurons for whole-cell patch clamp while stimulating ChR2-filled SNr axons [N=6]. (B) Percent connected among patched neurons in the rostral and caudal regions. (C) Average oIPSC amplitude at each of 40 optogenetic light pulses in n=9 rostral neurons and n=13 caudal neurons. (D) *Left*, Individual cell data for the first oIPSC amplitude and, *right*, example current traces. (E) Cell mapping of patched neuron locations with the first oIPSC amplitude represented by the color scale. (F) Normalized current amplitudes in C. (G) *Left*, Individual cell data for the PPR between the first two oIPSC amplitudes in the train and, *right*, example current traces. (H) Percent of pre-optical stimulation firing frequency [% Pre-Opto Frq] during stimulation and and rebound in n=7 rostral and n=19 caudal neurons. (I) Individual cell data for the absolute change in frequency during optical stimulation [ΔFrq During Opto]. (J) Correlation analysis, color scale representing Spearman r [-1,1] and size representing p-value [1,0]. (K) Negative correlation between the absolute change in frequency during stimulation and first oIPSC amplitude; r=-0.755, p=0.001. (L) Negative correlation between the pre-optical stimulation firing frequency and the PPR; r =-0.706, p=0.002. Box plots show median line with boxes showing IQR and whiskers showing 9th and 91st percentiles.

Although there was no major difference in the proportion of GABAergic neurons receiving inhibitory input from the SNr in the rostral vs caudal PPN, we compared the current amplitudes to characterize the synaptic strength and pattern of this connection. The oIPSC amplitudes across the 2-second stimulation as well as the first oIPSC showed no difference between rostral and caudal GABAergic neurons [median (IQR); first oIPSC n=9 Rostral: 76.5 pA (27.0 to 209.9 pA), n=13 Caudal: 59.7 pA (37.7 to 269.7 pA); Mann Whitney, U=50, p=0.6005; **Figure 3C&D**]. We also did not observe significant differences in short-term synaptic plasticity between rostral and caudal GABAergic neurons [median (IQR); PPR n=9 Rostral: 1.01 (0.88 to 1.16), n=13 Caudal: 1.06 (0.96 to 1.15); Mann Whitney, U=48, p=0.5123; **Figure 3 F&G**]. Defining short-term depression as a PPR<0.95 and short-term potentiation as PPR>1.05, we found 3/9 rostral and 7/13 caudal GABAergic PPN neurons showed short-term potentiation, while 3/9 rostral and 3/13 caudal GABAergic PPN neurons showed short-term depression. These data suggest additional heterogeneity within the GABAergic PPN subpopulation. Similar to the SNr connection with cholinergic PPN neurons, SNr input to the GABAergic PPN does not result in significant short-term synaptic plasticity and does not differ in amplitude or characteristics between rostral and caudal regions.

Because GABAergic PPN neurons display spontaneous firing^50^, we recorded action potential firing in current clamp while optically stimulating the SNr axons. The percent inhibition was not significantly different between rostral and caudal neurons (mean ± SEM; %Frequency n=7 Rostral: During Stimulation 37.91 ± 15.91 %, Post Stimulation 100.0 ± 14.57 %; n=19 Caudal: During Stimulation 30.68 ± 8.03 %, Post Stimulation 113.2 ± 5.187 %; 2way ANOVA, Stimulation_F(1, 24)=44.63_ p<0.0001, Region_F(1, 24)=0.09642_ p=0.7588, Interaction_F(1, 24)=0.8841_ p=0.3565; **Figure 3H**). There was also no difference in the absolute frequency change during optical stimulation between rostral and caudal neurons [median (IQR); ΔFrq During Opto n=7 Rostral: −5.03 Hz (−5.55 to −1.34 Hz), n=19 Caudal: −4.29 Hz (−11.02 to −2.94 Hz); Mann Whitney, U=55, p=0.5336; **Figure 3I**]. These data show that the SNr equally inhibits action potential firing in rostral and caudal GABAergic PPN neurons and does not evoke significant rebound firing in either neural population.

Overall, we found that the SNr inhibits the rostral and caudal GABAergic PPN neurons to a similar extent, but observed greater variability in synaptic strength and impact on neuronal action potential firing among caudal GABAergic neurons. One caudal subgroup responded with very small (<60 pA) amplitude currents and another subgroup responded with currents more than triple that amplitude, with one neuron responding with a 2.2 nA inhibitory current. We mapped the location of each recorded GABAergic neuron, as described above, and found no specific location corresponding with the neurons receiving stronger SNr inhibition across the PPN landscape (**Figure 3E**). This strong variability suggests that GABAergic PPN neurons may be divided into functionally distinct neural types that do not correspond to their rostrocaudal anatomical location. To determine if the neurons receiving larger inhibitory currents have distinct characteristics, we performed a correlation analysis (**Figure 3J**). As expected, we found that the absolute frequency reduction during inhibition is correlated with the first oIPSC amplitude recorded in a cell (Spearman r=-0.755, p=0.001; **Figure 3K**). Surprisingly, however, we found a significant negative correlation between PPR and the pre-optical stimulation frequency (Pre-Opto Frq, **Figure 3L**). Specifically, input to slower-firing neurons displayed greater short-term synaptic facilitation. This finding supports the idea that multiple functional neural subpopulations may be present within the GABAergic PPN population. Together, these results show that the SNr functionally inhibits GABAergic neurons across the rostrocaudal extent of the PPN, but suggests the presence of additional heterogeneity within the GABAergic PPN population.

### The SNr differentially inhibits the rostral and caudal glutamatergic PPN neurons

Previous rabies tracing anatomical studies have identified SNr inputs to glutamatergic PPN neurons^14,16,17^, but a full characterization of their synaptic strength and rostral and caudal connectivity has not been conducted. In Vglut2-Cre/Ai9-tdTomato mice, we injected ChR2 into the SNr to optically stimulate axons over the PPN while patching Vglut2+ glutamatergic neurons in the PPN (**Figure 4A**). While holding cells at −50mV and applying 470 nm blue light stimulation, neurons were identified as connected if oIPSCs were observed. Similar to our findings in cholinergic and GABAergic neurons, we found that all rostral (n=19/19, N=6) and all caudal (n=28/28, N=6) recorded glutamatergic neurons received inhibitory input from the SNr (**Figure 4B**).

**Figure 4.**
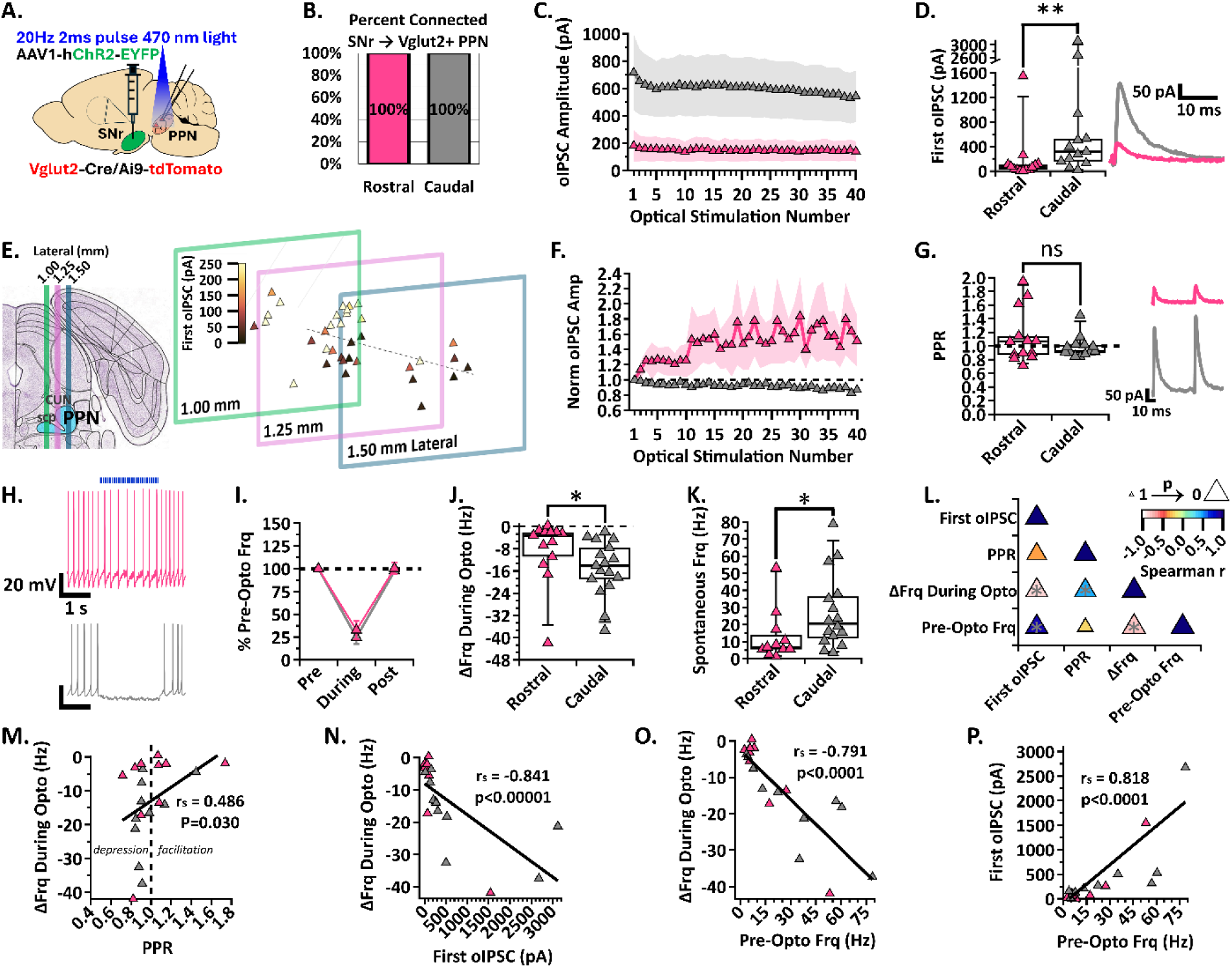
SNr inhibition of rostral and caudal Vglut2+ PPN neurons. (A) Experimental set up to identify red Vglut2+ PPN neurons for whole-cell patch clamp while stimulating ChR2-filled SNr axons [N=6]. (B) Percent connected among patched neurons in the rostral and caudal regions. (C) Average oIPSC amplitude at each of 40 optogenetic light pulses in n=13 rostral neurons and n=13 caudal neurons. (D) *Left*, Individual cell data for the first oIPSC amplitude and, *right*, example current traces; p=0.0035. (E) Cell mapping of patched locations with the first oIPSC amplitude represented by the color scale. (F) Normalized current amplitudes in C. (G) *Left*, Individual cell data for the PPR between the first two oIPSC amplitudes in the train and, *right*, example current traces. (H) Example voltage traces of action potential firing during a 2-second 20 Hz train stimulation in rostral and caudal neurons, top to bottom. (I) Percent of pre-optical stimulation firing frequency [% Pre-Opto Frq] during stimulation and rebound in n=13 rostral and n=17 caudal neurons. (J) Individual cell data for the absolute change in frequency during optical stimulation [ΔFrq During Opto]; p=0.0197. (K) Spontaneous frequency in n=11 rostral and n=16 caudal neurons; p=0.0343. (L) Correlation analysis, color scale representing Spearman r [-1,1] and size representing p-value [1,0]. (M) Positive correlation between the absolute change in frequency during stimulation and PPR, r=0.486, p=0.030. (N) Negative correlation between the absolute change in frequency during stimulation and first oIPSC amplitude; r=-0.841, p<0.00001. (O) Negative correlation between the absolute change in frequency during stimulation and pre-optical stimulation frequency; r=-0.791, p<0.0001. (P) Positive correlation between the first oIPSC amplitude and pre-optical stimulation frequency; r=0.818, p<0.0001. * p<0.05, ** p<0.01; box plots show median line with boxes showing IQR and whiskers showing 9th and 91st percentiles.

Comparing the oIPSC amplitudes across the 20 Hz train, we discovered that the caudal glutamatergic neurons receive larger inhibitory currents than rostral glutamatergic neurons (**Figure 4C**). The median first oIPSC measured in caudal glutamatergic neurons was also significantly larger than those recorded in rostral neurons [median (IQR); first oIPSC n=13 Rostral: 61.6 pA (19.2 to 113.9 pA), n=13 Caudal: 313.5 pA (143.8 to 729.1 pA); Mann Whitney, U=29, p=0.0035; **Figure 4D**]. After post-hoc staining for the cholinergic PPN and mapping the location of each recorded glutamatergic PPN neuron as described above, we identified a ‘hotspot’ of strong SNr inhibition in a medial-caudal group of glutamatergic neurons (**Figure 4E**). This finding shows that a subset of caudal PPN neurons preferentially receive exceptionally strong SNr input.

To determine whether the SNr input to the rostral and caudal PPN show differential short-term synaptic plasticity, we compared the normalized oIPSC amplitudes. We found that inhibitory inputs to the rostral glutamatergic neurons display short-term synaptic facilitation with amplitudes increasing in subsequent stimulations (**Figure 4F**). However, strong facilitation occurred in only a few neurons that drove the mean upward and there was no significant difference between the PPR observed in rostral and caudal glutamatergic neurons when activating SNr axons [median (IQR); PPR n=13 Rostral: 1.07 (0.86 to 1.38), n=13 Caudal: 0.92 (0.89 to 1.02); Mann Whitney, U=77, p=0.7241; Figure 4G].

Because glutamatergic PPN neurons spontaneously fire^16,50^, we wanted to determine if larger inhibitory currents mediated larger decreases in the firing rate of caudal PPN neurons. Therefore, we recorded tonic action potential firing in glutamatergic PPN neurons while optically stimulating SNr axons (**Figure 4H**). Although inhibition measured as the percent of pre-optical stimulation firing frequency was not significantly different between rostral and caudal neurons (mean ± SEM; %Frequency n=13 Rostral: During Stimulation 32.62 ± 10.21 %, Post Stimulation 100.3 ± 6.113 %; n=17 Caudal: During Stimulation 24.71 ± 7.30 %, Post Stimulation 97.48 ± 2.228 %; 2way ANOVA, Stimulation_F(1, 28)=133.6_ p<0.0001, Region_F(1, 28)=0.5214_ p=0.4762, Interaction_F(1, 28)=0.1765_ p=0.6776; **Figure 4I**), we found that the absolute frequency decrease was significantly larger in caudal glutamatergic neurons compared to rostral neurons during SNr stimulation [median (IQR); ΔFrq During Opto n=13 Rostral: −3.21 Hz (−12.21 to −1.95 Hz), n=17 Caudal: −14.08 Hz (−20.04 to −6.07 Hz); Mann Whitney, U=55, p=0.0197; **Figure 4J**]. To examine the discrepancy between the percent decrease and the absolute decrease in firing frequency, we compared the spontaneous firing frequency between rostral and caudal glutamatergic neurons in cells that had no holding current applied. We found that caudal glutamatergic neurons fire faster than rostral glutamatergic neurons [median (IQR); Spontaneous Frequency n=11 Rostral: 6.8 Hz (5.4 to 17.2 Hz), n=16 Caudal: 20.6 Hz (11.2 to 37.2 Hz); Mann Whitney, U=45, p=0.0343; **Figure 4K**]. Together, these findings show that fast-firing caudal glutamatergic PPN neurons receive larger SNr-mediated IPSCs and display greater decreases in their action potential firing.

Since we found that the caudal glutamatergic neurons receive larger inhibitory currents and display greater decreases in firing rate during SNr axon stimulation, we performed a correlation analysis to evaluate the strength of the relationship among the synaptic and firing characteristics (**Figure 4L**). Interestingly, there was a weak but significant positive correlation between the PPR and frequency change during stimulation showing that input displaying short-term synaptic depression occurs in neurons with greater decreases in firing frequency (Spearman r=0.486, p=0.030; **Figure 4M**). As expected, SNr-mediated oIPSCs with larger amplitudes are correlated with greater decreases in firing frequency during stimulation (Spearman r=-0.841, p<0.00001; **Figure 4N**). Greater pre-optical stimulation firing frequency is correlated with larger frequency decreases during stimulation (Spearman r=-0.791, p<0.0001; **Figure 4O**). This finding is expected because the fast-firing neurons can be more strongly inhibited in terms of absolute decrease in action potential frequency (i.e. a floor effect). Surprisingly, however, pre-optical stimulation firing frequency is correlated with the first oIPSC amplitude (Spearman r=0.818, p<0.0001; **Figure 4P**), indicating that the SNr is more strongly connected to glutamatergic PPN neurons with faster firing rates.

These correlations highlight strong selectivity in SNr projections to PPN glutamatergic neurons. Overall, these findings support that the SNr selectively targets and strongly inhibits fast-firing caudal glutamatergic PPN neurons compared to rostral neurons, and that there is a medial-caudal hotspot where SNr inhibition of PPN glutamatergic neurons is particularly strong.

### The SNr most strongly inhibits caudal glutamatergic PPN neurons

Most of the previous PPN circuit work has been done with rabies tracing. Aside from technical limitations, rabies tracing can only indicate the presence of synaptic connections. Our electrophysiological findings comprehensively characterize the strength and characteristics of the SNr input to each of the PPN cell types in both the rostral and caudal regions. Using this comprehensive dataset, we compared SNr input across the different PPN cell types in each region under the same electrophysiological conditions. We found that the median amplitude of the first SNr-mediated oIPSC was the same among rostral PPN neurons irrespective of cell type [median (IQR); first oIPSC n=15 ChAT+: 74.7 pA (44.5 to 136.5 pA), n=9 Vgat+: 76.5 pA (27.0 to 209.9 pA), n=13 Vglut2+: 61.6 pA (19.2 to 113.9 pA); Kruskal-Wallis, H_(2,34)_=0.7002 p=0.7046; **Figure 5A**].

**Figure 5.**
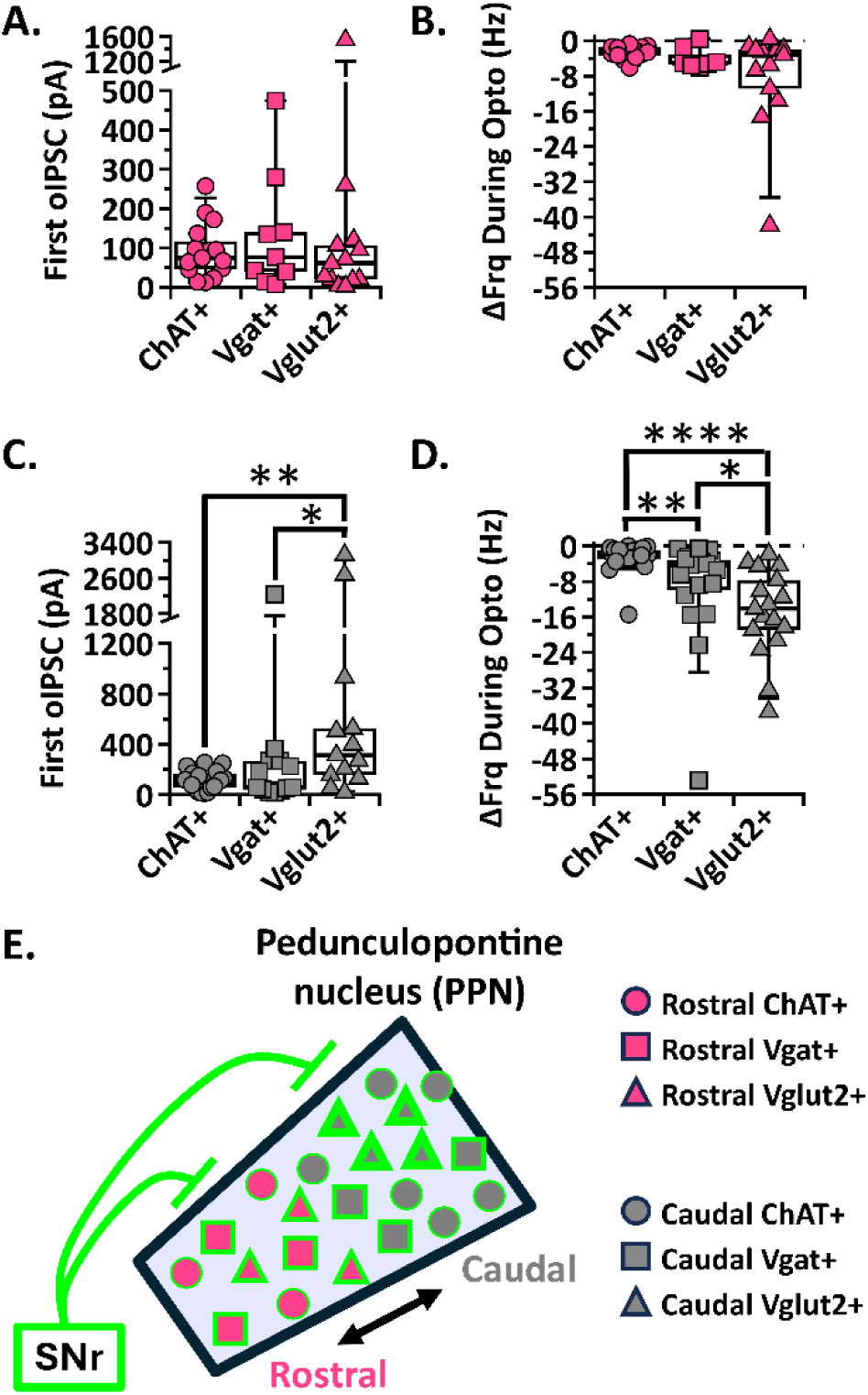
The SNr most strongly inhibits caudal glutamatergic PPN neurons. (A,C) Individual cell data for the first oIPSC amplitude recorded in each cell type for rostral and caudal PPN neurons, respectively. (B,D) Individual cell data for the absolute change in frequency during stimulation in each cell type for rostral and caudal PPN neurons, respectively. (E) Graphical depiction of SNr stimulation results. * p<0.05, ** p<0.01, **** p<0.0001; box plots show median line with boxes showing IQR and whiskers showing 9th and 91st percentiles.

The impact of these inhibitory inputs on neuronal firing did not differ with cell type as shown by the absolute change in frequency during SNr stimulation [median (IQR); ΔFrq During Opto n=14 ChAT+: −2.47 Hz (−3.37 to −1.44 Hz), n=7 Vgat+: −5.03 Hz (−5.54 to −1.34 Hz), n=13 Vglut2+: −3.21 Hz (−12.21 to −1.95 Hz); Kruskal-Wallis, H_(2,31)_=2.389 p=0.3028; **Figure 5B]**.

We found that the SNr-mediated oIPSCs in caudal glutamatergic neurons were significantly larger than in cholinergic and GABAergic caudal PPN neurons [median (IQR); first oIPSC n=20 ChAT+: 109.3 pA (58.5 to 168.6 pA), n=12 Vgat+: 59.7 pA (37.7 to 269.7 pA), n=13 Vglut2+: 313.5 pA (143.8 to 729.1 pA); Kruskal-Wallis, H_(2,43)_=8.717 p=0.0128; Dunn’s test, ChAT+ vs Vgat+ p=0.6409, ChAT+ vs Vglut2+ p=0.0042, Vgat+ vs Vglut2+ p=0.0295; **Figure 5C**].

Likewise, the impact of SNr inhibition on neuron firing rate was strongest in caudal glutamatergic neurons. The absolute frequency decrease during stimulation in caudal glutamatergic neurons was larger than both cholinergic and GABAergic neurons [median (IQR); ΔFrq During Opto n=23 ChAT+: −2.10 Hz (−3.11 to −0.84 Hz), n=19 Vgat+: −4.29 Hz (−11.02 to −2.94 Hz), n=17 Vglut2+: −14.08 Hz (−20.04 to −6.07 Hz); Kruskal-Wallis, H_(2,56)_=24.99 p<0.0001; Dunn’s test, ChAT+ vs Vgat+ p=0.0049, ChAT+ vs Vglut2+ p<0.0001, Vgat+ vs Vglut2+ p=0.0332; **Figure 5D**]. While we found no significant difference between SNr-mediated oIPSCs recorded in GABAergic and cholinergic neurons (**Figure 5C**), the GABAergic neuron firing frequency was more strongly inhibited than cholinergic (**Figure 5D**). The greater reduction in firing frequency may be driven by the caudal GABAergic subgroup receiving particularly large oIPSCs (**Figure 3D**) and could be explained if GABAergic PPN neurons have higher input resistance than cholinergic neurons, as has been previously suggested^59^. Altogether, our findings show that the SNr most strongly inhibits the glutamatergic neurons followed by GABAergic neurons in the caudal PPN and more weakly inhibits rostral PPN neurons without any cell type bias (**Figure 5E**).

### The GPe preferentially, but weakly, inhibits a subset of caudal GABAergic and glutamatergic PPN neurons

GPe projections to the cholinergic and glutamatergic PPN neurons have been shown in previous rabies tracing studies^14–17^, but projections to the GABAergic MLR neurons was not detected^14^. To evaluate the synaptic strength of these connections and determine regional connectivity, we repeated the previous whole-cell patch clamp experiments paired with optogenetics, but this time injecting ChR2 into the GPe of ChAT-Cre/Ai9-tdTomato, Vgat-Cre/Ai9-tdTomato, and Vglut2-Cre/tdTomato mice (**Figure 6A**). If oIPSCs were observed in response to optical stimulation, the neuron was identified as connected. In a subset of neurons, a GABA-a receptor blocker, GABAzine, was applied to confirm the oIPSCs were GABA-mediated (See *Methods* for details; **Figure 6B**). We recorded GPe-mediated oIPSCs in all three cell types of the PPN. As predicted by the low density of GPe axons present in the rostral PPN (**Figure 1D**), very few rostral PPN neurons received synaptic input from the GPe. This was true within each molecularly-defined PPN cell type [caudal vs rostral % connected; ChAT+: 30% vs 0% (n=9/30 vs 0/7), Vgat+: 68% vs 38% (n=21/31 vs 6/16), Vglut2+: 69% vs 26% (n=24/35 vs 5/19); **Figure 6Ci-iii**]. These findings show that the GPe preferentially targets caudal PPN neurons compared to rostral neurons. Since so few rostral PPN neurons received GPe inhibition, we focused our characterization of GPe inhibition to the caudal PPN.

**Figure 6.**
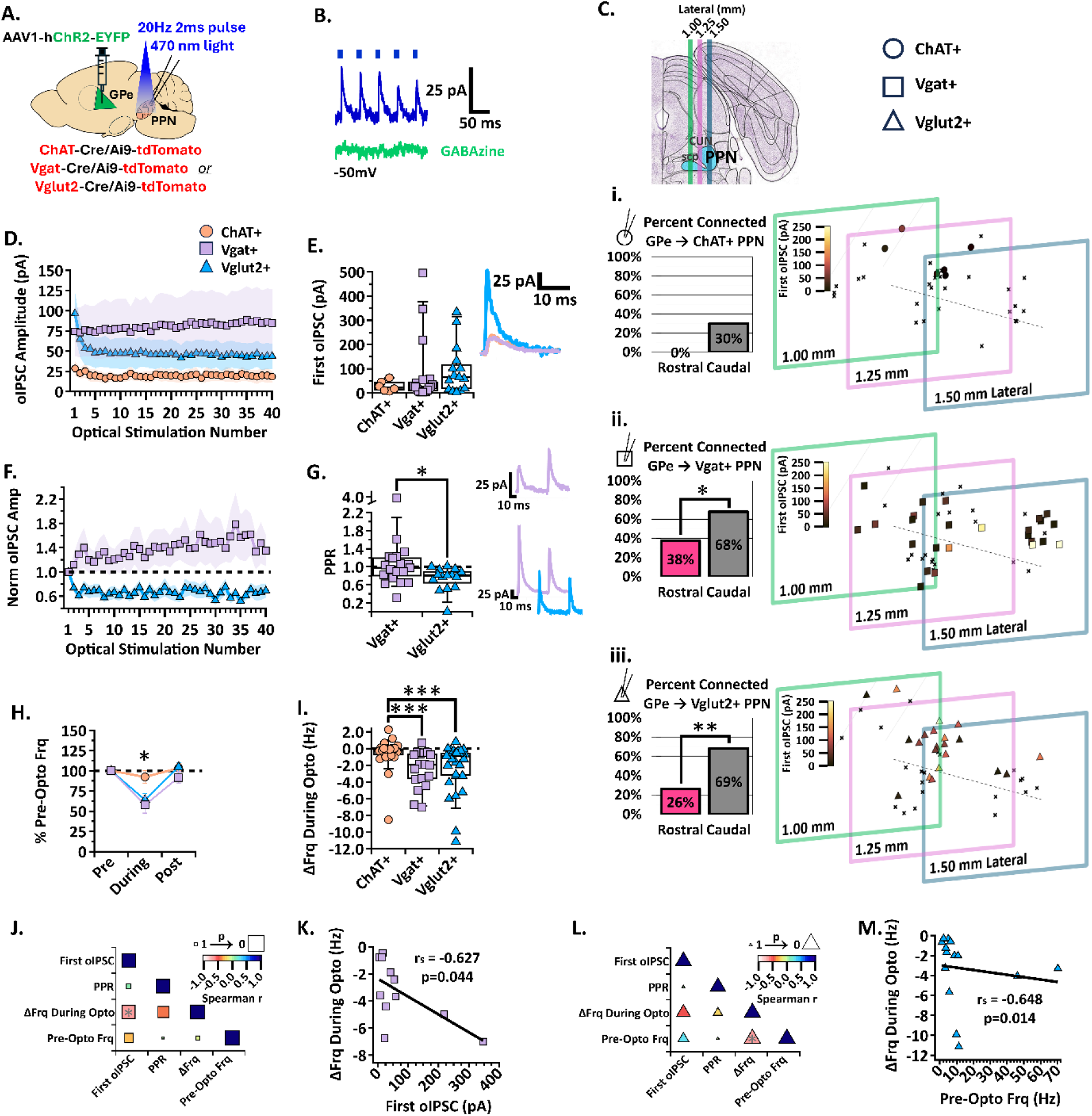
GPe inhibition of the three PPN cell types. (A) Experimental set up to identify red ChAT+, Vgat+, and Vglut2+ PPN neurons for whole-cell patch clamp while stimulating ChR2-filled GPe axons [N=6]. (B) Example trace of the first five oIPSCs in the 2-second 20 Hz train [blue] inhibited by GABA-a receptor blocker, GABAzine [green], while holding the cell at −50mV. (C) *Left*, Percent connected among patched neurons in the rostral and caudal regions and, *right*, cell mapping of patched locations with the first oIPSC amplitude represented by the color scale. *Top to bottom*, i. ChAT+, ii. Vgat+, and iii. Vglut2+ datasets. (D) Average oIPSC amplitude at each of 40 optogenetic light pulses in n=6 ChAT+, n=19 Vgat+, and n=15 Vglut2+ caudal PPN neurons. (E) *Left*, Individual cell data for the first oIPSC amplitude and, *right*, example current traces. (F) Normalized current amplitudes in C. (G) *Left*, Individual cell data for the PPR between the first two oIPSC amplitudes in the train; p=0.0206. R*ight, top*, example current trace of short-term synaptic facilitation in VgAT+ neurons. *Right, bottom*, example current traces of short-term synaptic depression in Vgat+ and Vglut2+ neurons. (H) Percent of pre-optical stimulation firing frequency [%Pre-Opto Frq] during and post-stimulation in n=25 ChAT+, n=18 Vgat+, and n=29 Vglut2+ caudal PPN neurons. (I) Individual cell data for the absolute change in frequency during stimulation [ΔFrq During Opto]. (J) Correlation analysis for Vgat+ neurons, color scale representing Spearman r [-1,1] and size representing p-value [1,0]. (K) Negative correlation between the absolute change in frequency during stimulation and first oIPSC amplitude; r=-0.627, p=0.044. (L) Correlation analysis for Vglut2+ neurons. (M) Negative correlation between the absolute change in frequency during stimulation and the pre-stimulation firing frequency; r=-0.648, p=0.014. *p<0.05, **p<0.01; box plots show median line with boxes showing IQR and whiskers showing 9th and 91st percentiles.

The percentage of connected caudal neurons was greater than the percentage of connected rostral neurons in both GABAergic and glutamatergic neurons (one-sided Fisher’s exact test; Vgat: OR=0.2857, CI=0.09190 to 0.9391, p=0.047; Vglut2: OR=0.1637, CI=0.05069 to 0.5767, p=0.0033; **Figure 6Cii&iii**), but we found no statistical difference between the rostral and caudal connectivity in cholinergic neurons (OR=0, CI=0 to 1.301, p=0.1150; **Figure 6Ci**). The number of connected caudal cholinergic neurons was not significantly greater than its 0% rostral connectivity (**Figure 6Ci**). In addition, the average oIPSC amplitude across the train and the first oIPSC measured in the connected cholinergic neurons was small [median (IQR); first oIPSC n=6 ChAT+: 22.6 pA (9.75 to 50.20 pA); **Figure 6D&E**]. Among recorded caudal cholinergic PPN neurons, both connected and non-connected, these small inhibitory currents displayed negligible effects on the neuronal firing frequency of the cholinergic PPN population in both the percent of pre-optical stimulation frequency and absolute frequency change [median (IQR); n=29 ChAT+: %Frequency During Stimulation 96.67 % (85.1 to 100.7 %); ΔFrq During Opto −0.16 Hz (−0.90 to 0.03 Hz); **Figure 6H&I**]. These findings indicate that the GPe has essentially no direct effect on the cholinergic PPN neural population. For this reason, we exclude GPe inhibition of cholinergic PPN neurons in the synaptic characterizations shown in Figures 6F&H. Together, we show that the GPe preferentially inhibits caudal GABAergic and glutamatergic neurons while largely avoiding rostral and cholinergic PPN neurons.

To determine whether the synaptic strength and characteristics from the GPe to the PPN differed between GABAergic and glutamatergic PPN subtypes, we evaluated the amplitude of the GPe-mediated oIPSCs during the 2-second 20 Hz optical stimulation. Both GABAergic and glutamatergic neurons responded to GPe stimulation with oIPSCs ranging from ones to hundreds of picoamperes (**Figure 6D**). While there was no significant difference in the first oIPSC measured in GABAergic and glutamatergic neurons [median (IQR); first oIPSC n=19 Vgat+: 24.77 pA (6.72 to 54.17 pA), n=15 Vglut2+: 65.71 pA (14.50 to 133.4 pA); Kruskal-Wallis, H_(2,37)_=3.050, p=0.2176; **Figure 6E**], the normalized amplitudes show that GPe input to GABAergic neurons facilitates with subsequent stimulations, while the input to glutamatergic neurons depresses (**Figure 6F**). The GPe-mediated inhibitory input to glutamatergic neurons displayed a significantly lower PPR than to GABAergic neurons [median (IQR); PPR n=19 Vgat+: 0.97 (0.76 to 1.2), n=15 Vglut2+: 0.81 (0.55 to 0.93); Mann-Whitney, U=76, p=0.0206; **Figure 6G**]. While short-term synaptic plasticity patterns were variable among GABAergic neurons with some depressing and others facilitating, GPe input to glutamatergic neurons generally depresses.

Short-term synaptic depression can dampen the impact that repeated IPSCs have on action potential output. Therefore, we compared the firing frequency change in each cell type during optogenetic stimulation of GPe axons. The PPN GABAergic neuron population firing rate was decreased to about 58% of their pre-stimulation firing with a median frequency decrease of 1.92 Hz during stimulation (**Figure 6H&I**). Similarly, the glutamatergic neuron population firing rate decreased to about 64% of their pre-stimulation firing with a median frequency decrease of 0.89 Hz during stimulation (**Figure 6H&I**). We found no statistical difference between GPe inhibition of firing frequency in caudal GABAergic and glutamatergic PPN neurons (mean ± SEM; %Frequency n=29 ChAT+: During Stimulation 92.39 ± 7.59 %, Post Stimulation 104.10 ± 5.23 %; n=18 Vgat+: During Stimulation 57.70 ± 10.11 %, Post Stimulation 91.81 ± 6.98 %; n=24 Vglut2+: During Stimulation 64.15 ± 7.63 %, Post Stimulation 104.7 ± 5.39 %; 2way ANOVA, Stimulation_F(1, 69)=25.46_ p<0.0001, Cell type_F(1, 69)=4.444_ p=0.0153, Interaction_F(1, 69)=2.805_ p=0.0674; Fisher’s LSD, df=138, %Frequency During Stimulation: ChAT+ vs. Vglut2+ p=0.0035, ChAT+ vs. Vgat+ p=0.0016, Vglut2+ vs. Vgat+ p=0.5378, %Frequency Post Stimulation: ChAT+ vs. Vglut2+ p=0.8883, ChAT+ vs. Vgat+ p=0.2838, Vglut2+ vs. Vgat+ p=0.2184; median (IQR); ΔFrq During Opto n=25 ChAT+: −0.16 Hz (−0.90 to 0.03 Hz), n=18 Vgat+: −1.92 Hz (−3.85 to −0.60 Hz), n=29 Vglut2+: −0.89 Hz (−3.28 to −0.43 Hz); Kruskall-Wallis, H_(2,69)_=17.21, p=0.0002; Dunn’s test, ChAT+ vs Vgat+ p=0.0002, ChAT+ vs Vglut2+ p=0.0009, Vgat vs Vglut2+ p=0.4020; **Figure 6H&I**). These findings show that the GPe inhibits GABAergic and glutamatergic neural populations to a similar extent.

To determine if there are significant relationships among the synaptic and firing characteristics, we performed a correlation analysis. Among GABAergic neurons receiving GPe inhibition, we found that the amplitude of the first oIPSC is significantly correlated with the absolute change in frequency during stimulation (Spearman r=-0.627, p=0.044, **Figure 6K**). Although GPe input to GABAergic neurons exhibited various PPRs, the correlation analyses revealed no significant relationship between PPR and absolute change in frequency during stimulation (**Figure 6J**). These findings suggest that the initial oIPSC amplitude of GPe input to GABAergic PPN neurons has a stronger relationship with firing rate than short-term synaptic plasticity does.

As nearly all GPe inputs to glutamatergic neurons exhibited short-term synaptic depression, we expected the correlation analysis to reveal a significant relationship between PPR and the change in firing frequency. However, similar to the GPe input to GABAergic neurons, PPR was not predictive of impact on neuronal activity (**Figure 6L**). GPe inhibition of the glutamatergic neurons showed a significant correlation between the absolute change in frequency during stimulation and pre-stimulation firing frequency (Spearman r=-0.648, p=0.014, **Figure 6M**). However, because the inhibitory synaptic current amplitude did not correlate with firing frequency, this correlation is likely due to a floor effect in which faster firing neurons have a greater capacity for reductions in firing frequency. Together, these findings show that the GPe preferentially inhibits caudal GABAergic and glutamatergic PPN neurons.

### *In vivo* activation of GPe and SNr axons in the PPN show differential effects on locomotion and valence

Once we established that the SNr and GPe inhibit different profiles of regionally- and molecularly-defined subpopulations of the PPN using *ex vivo* optogenetic electrophysiology, we wanted to evaluate the behavioral consequences of selectively stimulating these inputs *in vivo*. Direct optogenetic activation of PPN subpopulations have been previously shown to either promote or inhibit motion, sometimes with contradictory results^14,16,17,22–28^. To determine whether differential inhibitory inputs onto the PPN influenced locomotor activity, we bilaterally injected wild-type mice with ChR2 in either the SNr or GPe and implanted an optical fiber over the PPN (**Figure 7A&B**).

**Figure 7.**
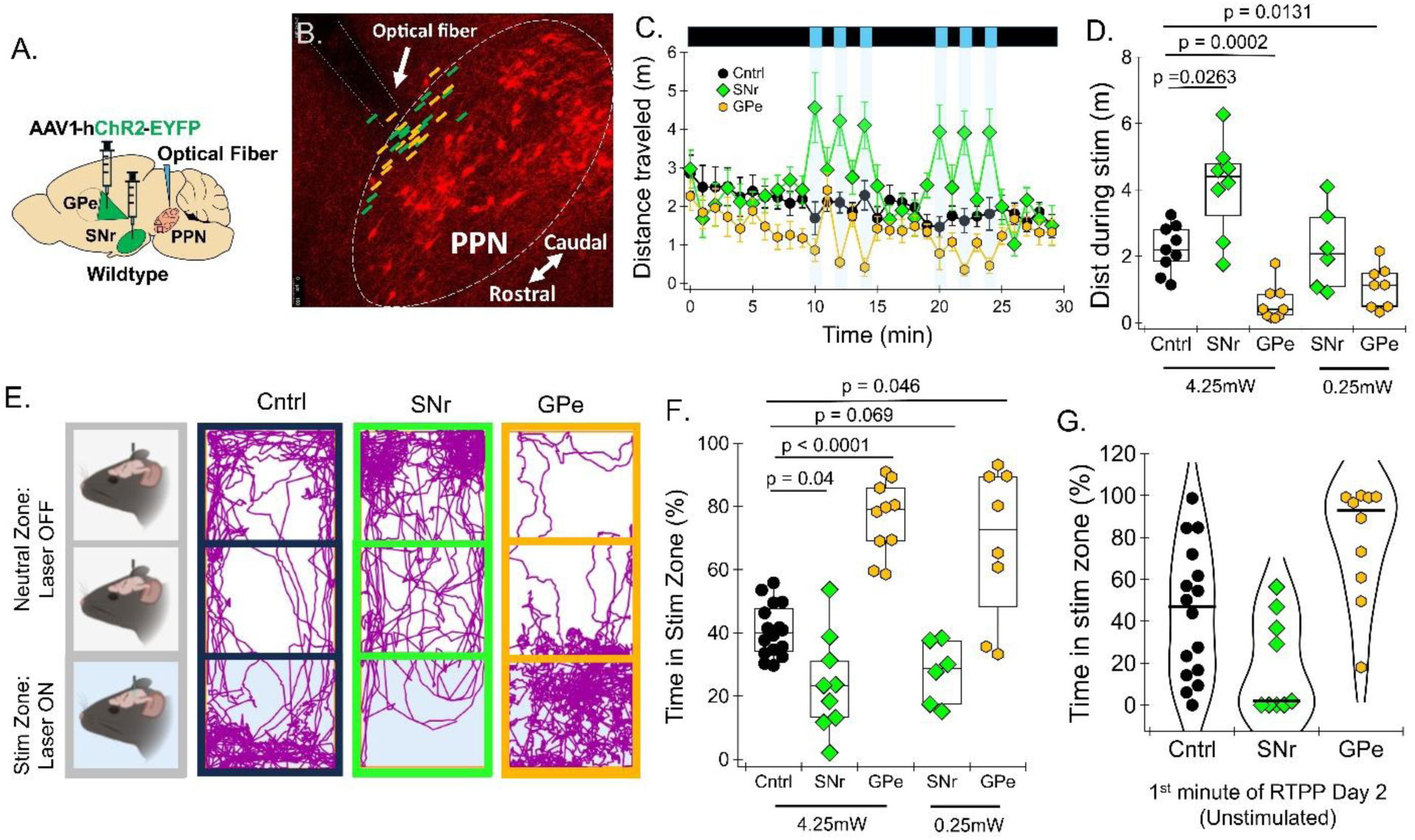
***In vivo* activation of GPe and SNr axons in the PPN show differential effects on ocomotion and opposite effects on valence.** (A) Experimental set up to stimulate ChR2-filled SNr or GPe axons over the PPN *in vivo*. (B) Representative image of optical fiber tract overlaid with the approximate optical fiber placement for SNr-[green] and GPe-[orange] injected mice. (C) Distance traveled over time in an open field with 1 minute 20 Hz 4.25 mW optical stimulations over the PPN in N=9 control (Ctrl) mice (black circles), N=8 mice injected with ChR2 in the SNr (green diamonds), and N=9 mice injected with ChR2 in the GPe (orange hexagons); vertical blue lines represent periods of optical timulation. (D) Average distance traveled for each mouse across the six 1-minute optical stimulations in the high (4.25 mW) and low laser power (0.25 mW) conditions (E) Representative mouse track tracings during real time place preference task in a three-chamber box and continuously stimulating EGFP- or ChR2-filled axons over the PPN at 20 Hz in SNr- and GPe-injected mice when the mice are in the stimulation zone. (F) Percent time spent in the stimulation zone in N=16 control mice, N=9 mice injected with ChR2 in the SNr, and N=10 mice injected with ChR2 in the GPe with 4.25 or 0.25 mW laser power. G) Percent time spent in the stimulation zone during the first minute reintroduced to the RTPP box on ay 2 of RTPP with the laser off (no chamber is stimulated) Black line = median. * p<0.05, ** p<0.01, *** <0.001, **** p<0.0001; box plots show median line with boxes showing IQR and whiskers showing 9th and 91st percentiles. See related supplemental Figures S2-S4 and Videos S1-S4.

Control mice were injected with an eGFP virus and implanted. After three weeks, we placed each mouse in the open field to measure gross locomotor behavior with and without optical stimulation of the SNr or GPe axons. We tracked the distance traveled for each mouse at baseline and during bilateral stimulation of either the SNr or GPe by applying blue light (473 nm, 4-4.5 mW) at 20 Hz with 2 ms pulses (4% duty cycle), for 1 minute at a time (**Figure 7C**). We found that stimulation of these two basal ganglia inputs to the PPN resulted in differential motor behaviors. Specifically, stimulation of SNr axons in the PPN increased distance traveled, while stimulation of GPe axons in the PPN decreased distance traveled [mean ± SEM; N=9 (3M, 6F) Control (Ctrl): 2.22 ± 0.24 m, N=8 (5M, 3F) SNr: 4.11 ± 0.51 m, N=9 (4M, 5F) GPe: 0.57 ± 0.18 m; Welch ANOVA p<0.0001; Dunnett’s test, Ctrl_4.25mW_ vs SNr_4.25mW_ p=0.0263, Ctrl_4.25mW_ vs GPe_4.25mW_ p=0.0002; **Figure 7D & Figure S2**, see Video S1&2].

The PPN has been implicated in reward processing and stimulation of the PPN can be reinforcing^7,28,29,31,35,60^. Therefore, we evaluated the effect of GPe or SNr axon stimulation in the PPN on valence in the real-time place preference (RTPP) task. In a three-chamber apparatus, the mouse could freely move between chambers for ten minutes. Optical stimulation of the SNr or GPe axons in the PPN was applied when the mouse entered the stimulated chamber and remained on at 20 Hz (4% duty cycle) until the mouse exited that chamber (**Figure 7E**). To reduce the motor effects of optical stimulation, this experiment was conducted with unilateral stimulation only. Interestingly, we again found opposite effects when stimulating the two basal ganglia inputs to the PPN. Mice avoided the stimulated chamber when the SNr axons were stimulated, but preferred the stimulated chamber when the GPe axons were stimulated (mean ± SEM; N=16 (6M, 10F) Ctrl: 40.76 ± 2.0 %, N=9 (5M, 4F) SNr: 23.99 ± 5.2 %, N=10 (5M, 5F) GPe: 76.11 ± 3.7 %; Welch ANOVA, p<0.0001; Dunnett’s test, Ctrl_4.25mW_ vs SNr_4.25mW_ p=0.0437, Ctrl_4.25mW_ vs GPe_4.25mW_ p<0.0001, Ctrl_4.25mW_ vs SNr_0.25mW_ p=0.0695, Ctrl_4.25mW_ vs GPe_0.25mW_ p=0.0467; **Figure 7F**, see Video S3&4). To further control for potential motion confounds, we recorded the location preference of the mice in the absence of optical stimulation the day after the stimulated RTPP assay (regardless of whether they received low or high power on day 1). The mice were placed in the central, neutral chamber and their location was tracked for 10 minutes. During the first minute of this assay, 8 out of 9 mice who had SNr optical stimulation on the first day continued avoiding the stimulated chamber while 9 out of 10 mice which had GPe optical stimulation continued to prefer the previously stimulated chamber compared to the median percent time spent in the stimulated chamber for mice in the control condition [median (IQR); N=16 (6M, 10F) Ctrl: 47% (14.79 to 69.24 %), N=9 (5M, 4F) SNr: 1.833% (0 to 41.75 %), N=10 (5M, 5F) GPe: 92.83% (58 to 99.21 %); Kruskall-Wallis, p=0.0008; Dunn’s test, Ctrl vs SNr p=0.1433, Ctrl vs GPe 0.0312; **Figure 7G**).

In addition to sending projections to the PPN, the GPe and SNr send axon collaterals to multiple other brain areas^12,37,61,62^. Therefore, it is possible that stimulating these axons in the PPN could cause antidromic stimulation of the GPe and SNr cell bodies, resulting in the inhibition of non-PPN brain regions. However, it has recently been shown that 0.25 mW laser power stimulation of axons can prevent antidromic activation of cell bodies^42^. Therefore, we repeated these optogenetic behavioral experiments using 0.25 mW laser power. We found that the locomotion effect of SNr stimulation over the PPN was no longer present during low power stimulation, but GPe stimulation still significantly decreased locomotion (mean ± SEM; distance traveled N=9 (3M, 6F) Ctrl: 2.22 ± 0.24 m, N=6 (3M, 3F) SNr: 2.24 ± 0.50 m, N=8 (4M, 4F) GPe: 1.09 ± 0.22 m; Welch ANOVA p<0.0001; Dunnett’s test, Ctrl_4.25mW_ vs SNr_0.25mW_ p>0.999, Ctrl_4.25mW_ vs GPe_0.25mW_ p=0.0131; **Figure 7D**). In the RTPP task, SNr axon stimulation continued to evoke place aversion at the trend level and GPe axon stimulation continued to significantly evoke place preference (mean ± SEM; % time in stimulation zone N=16 (6M, 10F) Ctrl: 40.76 ± 2.1 %, N=6 (3M, 3F) SNr: 27.72 ± 4.0 %, N=8 (4M, 4F) GPe: 68.41 ± 8.5 %; Welch ANOVA, p<0.0001; Dunnett’s test, Ctrl_4.25mW_ vs SNr_0.25mW_ p=0.0695, Ctrl_4.25mW_ vs GPe_0.25mW_ p=0.0467; **Figure 7F**). These findings show that decreased locomotion and place preference are mediated by GPe inhibition of the PPN while aversion is mediated by SNr inhibition of the PPN.

The increase in locomotion observed when stimulating the SNr axons in the PPN with high laser power is unexpected given that previous studies directly stimulating SNr neurons report decreases in movement^34,36^. Because injecting constitutive AAVs into the SNr can infect both GABAergic neurons and adjacent dopaminergic neurons of the substantia nigra pars compacta (SNc), we wanted to determine whether stimulating dopaminergic axons in the PPN enhanced locomotion and evoked aversion. To test this, we injected a cre-dependent ChR2 into the substantia nigra of DAT-cre mice (3M heterozygotes and 3F homozygotes) and ran open field and RTPP behavioral assays. We found that selective stimulation of dopaminergic axons in the PPN increased locomotion at high laser power (**Figure S3A**) and had no effect on locomotion at low power (**Figure S3C**). This finding suggests that dopaminergic neurons in the SNc send collateral projections to both the PPN and other locomotor-related structures (e.g. the striatum) such that stimulation of PPN-projecting SNc neurons increases locomotion.

The effect of dopaminergic axon stimulation did not differ from whole substantia nigra stimulation in either high (mean ± SEM; N=8 (5M, 3F) SNr_4.25mW_: 4.107 ± 0.51 m, N=6 (3M, 3F) DAT-Cre_4.25mW_: 3.778 ± 0.28 m; unpaired t-test, t=0.5159, df=12, p=0.6153; **Figure S3B**) or low laser power (mean ± SEM; N=6 (3M, 3F) SNr_0.25mW_: 2.238 ± 0.58 m, N=9 (3M, 3F) DAT-Cre_0.25mW_: 1.984 ± 0.30 m; unpaired t-test, t=0.4350, df=10, p=0.6728; **Figure S3C**). On the other hand, we found that stimulation of dopaminergic axons in the PPN did not cause place aversion, but rather induced real-time place preference at both high and low laser power (**Figure S3D-F**). Selectively stimulating dopaminergic axons induced place preference behavior significantly different from and opposite to whole substantia nigra stimulation at high (mean ± SEM; N=9 (5M, 4F) SNr_4.25mW_: 23.99 ± 5.18 %, N=6 (3M, S3F) DAT-Cre_4.25mW_: 70.83 ± 10.06 %; unpaired t-test, t=4.545, df=13, p=0.0005; **Figure S3E**) and low laser power (mean ± SEM; N=6 (3M, 3F) SNr_0.25mW_: 27.72 ± 4.0 %, N=6 (3M, 3F) DAT-Cre_0.25mW_: 50.82 ± 4.8 %; unpaired t-test, t=3.697, df=10, p=0.0041; **Figure S3F**). Together, these findings show that the the nigral increase in locomotion is not mediated through the PPN, and that dopaminergic input from the SNc to the PPN is rewarding while GABAergic input from the SNr to the PPN is aversive.

## Discussion

In this study, we comprehensively characterize the synaptic strength and impact on neuron action potential firing in molecularly- and regionally-defined PPN subpopulations while stimulating the SNr or GPe using *ex vivo* electrophysiology and optogenetics. We also stimulated the SNr or GPe axons in the PPN *in vivo* which revealed that the SNr and GPe evoke differential locomotion and valence processing behaviors in non-canonical basal ganglia circuits. Together, these findings show that distinct and selective inhibition of PPN subpopulations by the SNr and GPe can alter behavioral output.

## Region- and cell type-specific inputs to the PPN

The SNr and GPe have been previously shown to form monosynaptic synapses with PPN neurons using rabies viral tracing^12–17^. While SNr input to cholinergic and glutamatergic PPN neurons has been reproduced across different studies^13–17^, SNr inputs to GABAergic PPN neurons has only been inferred from studies of the entire mesencephalic locomotor region (MLR), consisting of both the PPN and cuneiform nucleus^14^. Several studies have used slice electrophysiology to show that the SNr inhibits PPN neurons generally^32^ or to compare SNr input to cholinergic and non-cholinergic PPN neurons^48,51,52^. However, a full characterization of the SNr input to regionally- and molecularly-defined PPN neurons in adult animals has been lacking. Using *ex vivo* whole-cell patch recordings, we were able to measure inhibitory currents in each of the PPN cell types during SNr stimulation showing that the SNr functionally inhibits each cell type within the rostral PPN to a similar extent while most strongly inhibiting a medial-caudal ‘hotspot’ of glutamatergic neurons.

While the GPe has been shown to project to the cholinergic and glutamatergic PPN neurons^15–17^, our electrophysiological results show that GPe inhibition of the cholinergic neurons is weak to non-existent and its inhibition of the glutamatergic PPN neurons is caudally biased. In contrast to a previous study showing that the GPe does not project to GABAergic MLR neurons^14^, we find that the GPe also preferentially inhibits a caudal subgroup of GABAergic PPN neurons. While the GPe selectively inhibits a caudal subpopulation of GABAergic and glutamatergic PPN neurons, its inhibition is much weaker than SNr inhibition of the PPN in terms of the proportion of connected PPN neurons, the inhibitory current amplitudes, and the inhibition of neuronal firing. Overall, our findings utilize a systematic approach to characterize the synaptic strength between these inhibitory basal ganglia nuclei and the PPN across its rostrocaudal axis for each cell type allowing us to compare SNr and GPe to PPN circuitry with regional-level granularity. These results also encourage future work involving the PPN to consider the individual influence of its rostral and caudal regions.

### Noncanonical GPe motor circuits

We found that stimulation of GPe axons over the PPN decreased locomotion in both low (putatively local) and high (putatively generating antidromic activity) laser power stimulations. This behavioral outcome appears counter to the canonical model of basal ganglia movement pathways, in which GPe activation increases movement. Although there is strong support for this canonical model, recent *in vivo* recordings have found subpopulations of GPe neurons that show activity patterns counter to this model (neurons that increase activity during immobility)^63,64^. Similarly, a subset of GPe neurons paradoxically increase activity upon indirect pathway striatal projection neuron stimulation^41^. While one explanation for these heterogeneous responses could be local inhibition within the GPe^40,41,65,66^, another possibility is that distinct subpopulations within the GPe differentially modulate locomotor behavior^39–41,61,67^. Our findings support non-canonical basal ganglia motor pathways involving GPe neurons that project to the PPN.

Our characterization of region-specific inhibitory inputs to the PPN helps us understand how GPe inhibition of the PPN can mediate decreases in locomotion. Our electrophysiology experiments show that the GPe exerts minimal influence on the rostral PPN neurons which appear to decrease locomotion^22,24,25^ and preferentially inhibits caudal GABAergic and glutamatergic PPN neurons. Because stimulating the caudal GABAergic and glutamatergic PPN neurons increases locomotion^23^, GPe selective inhibition of these caudal neurons could explain our behavioral experiments showing that stimulation of GPe axons over the PPN decreases locomotion. Future studies are needed to determine if GPe inhibition of both cell types is necessary for mediating decreased locomotion. Aligned with our findings, selectively stimulating the Npas1+ or FoxP2+ subpopulations in the GPe decreases locomotion^39,41^. Some groups have found that direct stimulation of the PV+ GPe neurons increases locomotion^39,62^; however, one study shows that inhibition of PV+ GPe neurons can increase locomotion^42^. While we show that GPe projections to the PPN decrease locomotion, future studies are needed to determine which GPe subpopulations project to the PPN to modulate locomotion.

### GABAergic vs dopaminergic effects of nigral projections to the PPN

While we found that stimulating nigral axons in the PPN with high laser power increased locomotion, we observed no locomotor effect when using low laser power stimulation. The simultaneous inhibition of caudal PPN neurons that promote movement^23^ and rostral PPN neurons that suppress movement^22,24,25^ may explain a net zero effect on locomotion during low power optical stimulation of SNr axons in the PPN. Indeed, low laser power stimulation of nigral axons in the most rostral implant location resulted in increased locomotion, while the most caudal implant location resulted in decreased locomotion (**Figure S4**). Interestingly, the enhanced locomotion observed during high laser power stimulation suggests that a subset of nigral neurons projecting to the PPN promote movement. However, the SNr canonically suppresses movement and the direct somatic stimulation of PV+ and GAD2+ SNr neurons that send axon collaterals to the PPN^12,68^ decreases gross locomotor movement^34,68^. Since injecting constitutive AAVs into the substantia nigra can infect both GABAergic neurons of the SNr and adjacent dopaminergic neurons of the SNc, it is possible that stimulation of dopaminergic activity may mask the impact of stimulating the GABAergic SNr alone.

Both the increased locomotion with high laser power and the lack of change with low laser power was recapitulated when we ran these experiments while selectively stimulating dopaminergic projections from the SNc to the PPN (**Figure S3**). Therefore, our findings suggest that activation of the PPN-projecting dopaminergic SNc neurons promotes locomotion, but not through direct actions at the PPN. This is consistent with previous literature showing that SNc dopaminergic neurons send axons to both the PPN and striatum^69^ and that direct stimulation of nigrostriatal dopaminergic neurons can increase movement initiation and vigor^70^. Therefore, antidromic stimulation of the SNc may evoke dopamine release into the striatum to increase movement in our high laser power experiments. Future studies are needed to evaluate the motor behavioral impact of the different SNr cell types when inhibiting the entire PPN versus the rostral or caudal PPN subregions.

By contrast, we find that both high and low power non-selective SNr axon stimulation in the PPN is aversive (**Figure 7E-G**), while both high and low power selective stimulation of dopaminergic SNc axons in the PPN is rewarding (**Figure S3**). These findings are particularly interesting because they indicate that the dopaminergic vs GABAergic inputs from the substantia nigra to the PPN serve opposite functions regarding valence signaling. Importantly, these findings indicate that GABAergic input from the SNr is highly aversive and that this aversive signaling occurs through the PPN. Interestingly, these findings also show that dopaminergic SNc neurons do not require antidromic stimulation to signal reward. This means that dopaminergic signaling within the PPN may serve to reinforce behavior. Future work is needed to parse out specific reward and aversion pathways involving neural subpopulations within the SNr, SNc, and PPN.

### Differential PPN inhibition by the SNr and GPe mediate opposing valence processing outcomes

We found that stimulation of SNr axons over the PPN was aversive in real time place preference (RTPP), while stimulation of GPe axons over the PPN led to a preference for the stimulated chamber. Previous studies directly stimulating SNr and GPe subpopulations have implicated these structures in reward and aversion processing.

Inhibition of Vgat+ SNr neurons has been shown to increase place preference^36^ while its direct stimulation has no effect^34,36^. However, selective stimulation of the PV+ SNr neurons induces aversion in RTPP^34^. Although the GPe has been predominantly studied in the context of movement, recent work has begun to reveal a role for the GPe in valence processing^42,71–73^. One recent study shows that inhibition of PV+ GPe neurons induced aversion while inhibition of Npas1+ GPe neurons induced placed preference in RTPP^42^. Because activation of PV+ SNr neurons is aversive^34^ and inhibition of PV+ GPe neurons is aversive^42^, it is likely that our results are due to activation of the PV+ SNr and GPe axons in the PPN. While we show that activation of the PPN-projecting SNr and GPe subpopulations strongly affect valence processing, future studies are needed to determine whether these effects are specifically mediated by PV+ SNr and GPe projections to the PPN.

Neurons in the PPN have been implicated in reward and aversive-related behaviors^7,28,29,31–33,35,60^. The cholinergic PPN neurons are implicated in positive valence with place preference and increased lever pressing through their projections to the ventral tegmental area (VTA)^28,29^. Aligned with our findings, direct inhibition of the cholinergic PPN neurons evokes place aversion^28^. Of particular significance, we show that the SNr inhibits the cholinergic PPN neurons while the GPe axons avoid cholinergic PPN neurons. Therefore, it is possible that the difference between the SNr and GPe effects on valence is due to the difference in their inhibition of PPN cholinergic neurons.

Direct stimulation of the glutamatergic PPN neurons has also been implicated in positive valence with increased reinforcement behavior^7^. Both the cholinergic and glutamatergic PPN neurons directly influence dopamine release in the striatum and dopaminergic neuron activity in the SNc and VTA^7,28,29,74–78^. Specifically, the caudal PPN neurons are thought to innervate the medial part of the SNc and the VTA^9,78^, two major reward pathway hubs. Our data show that the SNr most strongly inhibits the caudal glutamatergic PPN neurons. Therefore, strong SNr inhibition of these neurons would remove excitatory drive from the reward-related dopaminergic neurons. This pathway could also contribute to the aversive effect of SNr axon stimulation in the PPN. Future experiments are needed to test whether direct inhibition of the caudal glutamatergic PPN neurons receiving SNr input is sufficient to evoke aversion.

The pathways underlying the GPe-mediated rewarding effect are less clear and may indicate functional heterogeneity among caudal PPN neurons. We find that the GPe only inhibits a subset of caudal GABAergic and glutamatergic PPN neurons. It is possible that the subset of GABAergic PPN neurons targeted by the GPe disinhibits local cholinergic and glutamatergic neurons to increase excitation of the rewarding dopaminergic neuron pathways. Another possibility is that the specific GPe-inhibited PPN population may have particularly aversive-properties. For example, PPN neurons that excite amygdala nuclei evoke place avoidance^35,60^. In this case, GPe inhibition of these neurons could be rewarding by removing excitation of amygdala nuclei involved in negative valence processing. To understand the PPN circuitry involved in GPe-mediated place preference, future experimental work must determine whether inhibiting GPe-targeted GABAergic or glutamatergic PPN subpopulations alone is sufficient to evoke place preference.

## Conclusions

We have systematically characterized SNr and GPe inputs across the rostrocaudal axis of the PPN for each cell type – cholinergic, GABAergic, and glutamatergic. We show that the SNr inhibits nearly every PPN cell recorded with differential strength determined by cell type and anatomical biases while identifying a medial-caudal ‘hotspot’ of glutamatergic neurons most strongly inhibited by the SNr. In contrast, the GPe strikingly avoids the cholinergic PPN neurons and weakly, but selectively, inhibits a subpopulation of caudal GABAergic and glutamatergic neurons. We proposed that the differential inhibition of regionally-biased, mixed-cell type PPN subpopulations can alter behavioral outputs. By stimulating these inhibitory basal ganglia axons over the PPN, we show that the SNr evokes place aversion while the GPe evokes place preference and decreases locomotion. Together, our findings show that the SNr and GPe mediate opposing valence processing outcomes through the PPN and support a non-canonical basal ganglia motor pathway in which the GPe decreases locomotion through its projections to the PPN.

## Acknowledgements

Funding for this research was provided by an NIH BRAIN Initiative grant R00NS112417 (to RCE), NIH NINDS grant 1F30NS132399 (to MF), Michael J Fox Foundation Research Grant MJFF-022863 (to RCE), and Parkinson’s Foundation Summer Student Fellowship PF-SSF-941810 (to AES). This research was supported by the Microscopy and Imaging Shared Resource (MISR, S10RR025661) and the Lombardi Comprehensive Cancer Center grant (P30-CA051008). We would like to thank the Division of Comparative Medicine for their animal care and management and Dr. Stefano Vicini, Dr. Aryn Gittis, and members of the Evans lab for their feedback on earlier versions of this manuscript.

## Author Contributions

M.F. and R.C.E. conceptualized the experiments; M.F. performed stereotaxic injections, conducted the electrophysiological experiments, and analyzed the data; A.E.S. and K.C.U. performed stereotaxic injections, conducted the behavioral experiments, and analyzed the data; C.B.S. performed stereotaxic injections for pilot experiments; M.F. wrote the initial draft of the manuscript; M.F. and R.C.E. edited and revised the manuscript.

## Star Methods text

### KEY RESOURCES TABLE

| REAGENT or RESOURCE | SOURCE | IDENTIFIER |
| --- | --- | --- |
| **Antibodies** |  |  |
| goat anti-choline acetyltransferase (ChAT) | Millipore | Cat#: AB144P,<br>RRID: AB_2079751 |
| Streptavidin, Cy5 | Invitrogen | Cat#: SA1011 |
| Streptavidin, DyLight™ 405 | Invitrogen | Cat#: 21831 |
| Donkey anti-goat IgG (H+L) Cross-Adsorbed Secondary Antibody, Alexa Fluor™ 647 | Invitrogen | Cat#: A-21447,<br>RRID: AB_2535864 |
| Sheep anti-tyrosine hydroxylase (TH) | Novus Biologicals | Cat# NB300-110,<br>RRID: AB_10002491 |
| DyLight™ 405 AffiniPure™ Donkey Anti-Sheep IgG (H+L) | Jackson ImmunoResearch Laboratories | Cat# 713-475-147,<br>RRID: AB_2340740 |
| **Bacterial and virus strains** |  |  |
| AAV1-hSyn-hChR2(H134R)-EYFP | Addgene (Deisseroth Lab) | Cat#: 26973 |
| AAV1-Ef1a-double floxed-hChR2(H134R)-EYFP-WPRE-HGHpA | Addgene (Deisseroth Lab) | Cat#: 20298 |
| AAV1-hSyn-EGFP | Addgene (Roth Lab) | Cat#: 50465 |
| **Chemicals, peptides, and recombinant proteins** |  |  |
| D-AP5 | Tocris and HelloBio | Cat#: 0106 and<br>HB0225 |
| NBQX | Tocris and HelloBio | Cat#: 1044 and<br>HB0443 |
| CNQX disodium salt | HelloBio | Cat#: HB0205 |
| SR 95531 hydrobromide (GABA<sub>A</sub>zine) | Tocris and HelloBio | Cat#: 1262 and<br>HB0901 |
| **Experimental models: Organisms/strains** |  |  |
| ChAT-cre mice: B6.129S-Chat<sup>tm1(cre)Lowl</sup>/MwarJ | Jackson | RRID:<br>IMSR_JAX:031661 |
| Vgat-cre mice: B6J.129S6(FVB)-*Slc32a1*<sup>tm2(cre)Lowl</sup>/MwarJ | Jackson | RRID:<br>IMSR_JAX:028862 |
| Vglut2-cre mice: B6J.129S6(FVB)-*Slc17a6*<sup>tm2(cre)Lowl</sup>/MwarJ | Jackson | RRID:<br>IMSR_JAX:028863 |
| Ai9-tdTomato mice: B6.Cg-Gt(ROSA)26Sor<sup>tm9(CAG-tdTomato)Hze</sup>/J | Jackson | RRID:<br>IMSR_JAX:007909 |
| DAT-cre mice [B6.SJL-Slc6a3<sup>tm1.1(cre)Bkmn</sup>/J | Jackson | RRID:<br>IMSR_JAX:006660 |
| C57BL/6J | Jackson | RRID:<br>IMSR_JAX:000664 |
| **Software and algorithms** |  |  |
| Igor Pro | WaveMetrics | Version 9.00 |
| GraphPad Prism | Dotmatics | Version 9.5.0 |
| Fiji software | Open source on GitHub | Version 1.51n |
| ANY-maze® software | Stoelting Company, Wood Dale, IL. | Version 7.36 |
| Clampex software | Molecular Devices | Version 11.2 |
| **Other** |  |  |
| Multiclamp™ 700B amplifier |  |  |
| Axon™ Digidata® 1550B |  |  |
| Leica SP8AOBS++ |  |  |
| Zeiss Axio Imager Z2 |  |  |
| Sutter Instrument Model P-97 |  |  |

### RESOURCE AVAILABILITY

#### Lead contact

Further information and requests for resources and reagents should be directed to and will be fulfilled by the lead contact, Dr. Rebekah C. Evans.

#### Materials availability

This study did not generate new unique reagents.

#### Data and code availability

- All data and code supporting the current study will be shared by the lead contact upon request.
- No original code was generated in this paper.
- Any additional information required to reanalyze the data reported in this paper is available from the lead contact upon request.

### EXPERIMENTAL MODEL AND STUDY PARTICIPANT DETAILS

#### Animal Welfare

All animal procedures were approved by the Georgetown University Medical Center Institutional Animal Care and Use Committee (IACUC). Measures were taken to ensure minimal animal suffering and discomfort, and protocols were designed to minimize the number of animals used.

#### Animal Subjects

Homozygous Ai9-TdTomato mice [B6.Cg-Gt(ROSA)26Sor^tm9(CAG-tdTomato)Hze^/J; JAX# 007909] were bred with homozygous ChAT-Cre mice [B6.129S-Chat^tm1(cre)Lowl^/MwarJ; JAX# 031661], homozygous Vgat-Cre mice [B6J.129S6(FVB)-*Slc32a1^tm2(cre)Lowl^*/MwarJ; JAX# 028862], and Vglut2-Cre mice [B6J.129S6(FVB)-Slc17a6^tm2(cre)Lowl^/MwarJ; JAX# 028863] to obtain ChAT-Cre/Ai9-tdTomato, Vgat-Cre/Ai9-tdTomato, and Vglut2-Cre/Ai9-tdTomato in-house, respectively. C57BL/6J wildtype mice (Jax# 000664) and DAT-cre mice [B6.SJL-Slc6a3^tm1.1(cre)Bkmn^/J; JAX#006660] were obtained from Jackson for behavioral experiments. Animals were housed under a 12:12 light-dark cycle (lights on at 06:00 AM) with food and water *ad libitum*.

#### Experimental Groups

For electrophysiological studies, ChAT-Cre/Ai9-tdTomato, Vgat-Cre/Ai9-tdTomato, and Vglut2-Cre/Ai9-tdTomato mice of both sexes (3 males and 3 females in each group) aged 2-5 months were used. For behavioral studies, C57BL/6J wildtype mice of both sexes, with age-matched littermates randomly assigned to control or experimental groups. DAT-cre mice of both sexes were used for behavioral experiments where specified. Behavioral assessments were exclusively conducted during the light phase, and all testing chambers were cleaned with 70% ethanol between mice to mitigate potential olfactory influences.

### METHOD DETAILS

#### Viral injections and optical fiber implantation surgeries

Mice at least 7 weeks old were briefly anesthetized with inhaled 5% isoflurane using an anesthetic chamber and placed onto a heated pad within the stereotaxic frame (Stoelting 51730UD). The skull was stabilized with evenly positioned ear bars and nose cone properly positioned to deliver continuous 1-3% isoflurane and oxygen at a steady flow of 1L/min throughout the surgery duration. Bupivacaine (5 mg/kg) and carprofen (5mg/kg) were administered as local anesthetic and analgesic, respectively. A small incision was made on the scalp to visualize bregma and lambda, which were used as references to level the skull. Bilateral holes were drilled at either the coordinates of the SNr (AP −3.1 mm, ML ± 1.4 mm, DV − 4.7 mm relative to bregma) or GPe (AP −0.3 mm, ML ± 1.9mm, DV −3.9 mm relative to bregma). The 5µL Hamilton microsyringe was positioned and 250 nL virus was injected at a rate of 0.2 µL/min. The syringe was raised and rested 0.5 mm above the injection site for 10 minutes. For *ex vivo* optogenetic experiments, AAV1-hSyn-hChR2(H134R)-EYFP (2.3 x 10^13 particles per milliliter, Addgene, Cat# 26973) was injected into either the SNr or GPe. For *in vivo* optogenetic experiments, AAV1-hSyn-hChR2(H134R)-EYFP 2.0 x 10^13 particles per milliliter, Addgene, Cat #26973), AAV1-hSyn-EGFP (1.1 x 10^13 particles per milliliter, Addgene, Cat# 50465), or AAV1-Ef1a-double floxed-hChR2(H134R)-EYFP-WPRE-HGHpA (2.3 x 10^13 particles per milliliter, Addgene, Cat#20298) was injected into either the SNr or GPe for experimental and control mice, respectively. For mice used in the *in vivo* optogenetic experiments, an optical fiber (200 µm core, 0.22 NA, 3.7 mm length) was implanted over the middle-to-rostral PPN (AP −4.5, ML ± 1.1 mm, DV −3.5 relative to bregma). The skin on either side of the incision site was joined, glued together using VetBond tissue adhesive, and fastened with two wound clips or the optical fiber was fixed in position with dental cement. (C&B-Metabond Quick! Adhesive Luting Cement by Parkell Products Inc, Patterson# 553-3484/CAT# S380; Jet Denture Repair Package by LANG, CAT# 1223F2). Post-surgery, buprenorphine SR (1.5 mg/kg) was administered for long-acting analgesia. Mice were allowed to recover on a heating pad until fully awake and were monitored daily for signs of distress or infection.

#### Electrophysiological solutions

Acute brain slices were prepared in modified artificial cerebrospinal fluid (aCSF) solutions with final osmolarities ∼300-310 mOsm and ∼7.4 pH. The slicing solution contained (in mM) 198 glycerol, 2.5 KCl, 1.2 NaH_2_PO_4_, 20 HEPES, 25 NaHCO_3_, 10 glucose, 10 MgCl_2_, and 0.5 CaCl_2_. The holding solution contained (in mM) 92 NaCl, 2.5 KCl, 1.2 NaH_2_PO_4_, 30 NaHCO_3_, 20 HEPES, 2 MgCl_2_, 2 CaCl_2_, 35 glucose, 5 sodium ascorbate, 3 sodium pyruvate, and 2 thiourea. Recording aCSF was made up of (in mM): 125 NaCl, 25 NaHCO_3_, 3.5 KCl, 1.25 NaH_2_PO_4_, 10 glucose, 1 MgCl_2_, 2 CaCl_2_. Whole-cell patch clamp recordings used a potassium methane sulfonate (KMeSO_3_)-based internal solution containing (in mM) 122 methanesulfonic acid, 9 NaCl, 9 HEPES, 1.8 MgCl_2_, 4 Mg-ATP, 0.3 Tris-GTP, and 14 phosphocreatine for a final osmolarity between 290 and 305 mOsm. The internal solution contained neurobiotin (0.1-0.3%) for post-hoc staining. These reagents enable reliable electrophysiology recordings in adult brainstem neurons^46,79^.

#### Slicing and Electrophysiology

Animals ages 2-5 months old were anesthetized with inhaled isoflurane and transcardially perfused with ice-cold slicing solution that had been bubbled with 95% O_2_ and 5% CO_2_. Mice were decapitated and brains were quickly extracted from the skull, keeping the cerebellum intact. For sagittal slices, the brain hemispheres were separated and the medial side was glued onto a 3% agar block fixed to the stage of a semi-automatic Leica VT1200 microtome. 200 µm-thick slices were obtained and incubated at 34°C in holding solution for 30 minutes then kept at room temperature. In all steps, the modified aCSF solutions are bubbled with 95% O_2_ and 5% CO_2_. Slices in recording chamber during whole-cell patch clamp experiments were continuously perfused with oxygenated recording aCSF kept at 28-34°C using a water bath and in-line Warner heater. Cells were visualized using an Olympus OpenStand upright microscope and 565nm ThorLabs LED light. Recording pipettes with resistance between 1.5-4 MΩ were prepared using borosilicate glass capillaries (World Precision Instruments, Inc. 1B150F-4, 4-inch length, 1.5 mm OD, 0.84 mm ID) with a micropipette puller (Sutter Instrument Model P-97). Recordings were obtained using a Multiclamp^TM^ 700B amplifier and Axon^TM^ Digidata^®^ 1550B controlled by Clampex 11.2. Voltage-clamp signals were low-pass filtered at 2kHz and sampled at 10 kHz. Current-clamp signals were low-pass filtered at 10kHz and sampled at 10 kHz. At the end of recording, the cell was sealed by moving the recording pipette slightly above the soma and an image was taken of the PPN and pipette tip.

#### *Ex vivo* optogenetic activation

Whole-field optogenetic activation of channelrhodopsin-infected axons in brain slice was achieved with a blue (470 nm) ThorLabs LED light sent to the tissue via a silver mirror. Light intensity measured at the objective back aperture was 13 mW. Light activation was applied at 20 Hz with 2 ms pulse intervals for 2 seconds. All recordings were conducted in the presence of glutamatergic receptor blockers – 50µM D-AP5 (Tocris Cat#0106 and HelloBio Cat#H0225), and 5µM NBQX (Tocris Cat#1044 and HelloBio Cat#HB0443) alone or in combination with 20µM CNQX (HelloBio Cat#HB0205).

In voltage-clamp, cells were determined to be connected if observable optically evoked inhibitory post synaptic currents (oIPSCs) were measured while holding the cell at −50 mV. In a subset of neurons, 10µM GABAzine (Tocris Cat#1262 and HelloBio Cat#HB0901) was applied to ensure the oIPSCs were GABA-a receptor mediated. In a few cells (17/54), small inhibitory currents remained with amplitudes averaging about 24% of the first oIPSC measured. Any cells that required greater than 200pA to be held at −50mV or that had access resistance that exceeded 30 MΩ at the start of recording were excluded from oIPSC quantitative analyses as they were likely unhealthy and too leaky; however, the first oIPSC amplitude was used as an approximate measure of input strength in the cell maps (SNr: 21/46; GPe: 11/40). oIPSC amplitudes were measured from the baseline to the peak of each current. The paired pulse ratio (PPR) was calculated from the amplitudes of the first two currents evoked in the 20 Hz train.

For current-clamp recordings, most patched PPN neurons spontaneously fired, but a few required a small amount of hyperpolarizing or depolarizing current to fire [(R)ostral, (C)audal; SNr-PPN_ChAT_ R: 3/14, C: 4/23; SNr-PPN_Vgat_ R: 2/7, C: 9/19; SNr-PPN_Vglut2_ R: 2/13, C: 1/17; GPe-PPN_ChAT_ R: 2/7, C: 9/25; GPe-PPN_Vgat_ R: 2/7, C: 8/18; GPe-PPN_Vglut2_ R: 6/12, C: 7/29 required current). Neurons injected with more than 60 pA were excluded. In this manuscript, we use “pre-stimulation firing frequency” to refer to the time period right before optical stimulation. The pre-stimulation firing frequency was measured using the number of action potentials in the one-second epoch before the start of stimulation. The firing frequency during stimulation was measured using the middle one-second of the 2-second stimulation. The post-stimulation firing frequency was measured in the one-second epoch immediately following the end of the stimulation. The absolute frequency change is the difference between the pre-simulation and during stimulation frequencies. The rebound frequency change is the difference between the post-stimulation and pre-stimulation frequencies. These values include cells with and without holding current applied. By contrast, the spontaneous firing rate of the cell includes only cells with no holding current applied.

#### *In vivo* optogenetic activation

Two behavioral tests, the open field (OF) test and real-time place preference (RTPP), were conducted sequentially to assess gross locomotion and place preference in animals of at least 11 weeks old (mean age ± SEM; SNr-injected mice: 132 ± 23.4 days old; GPe-injected mice: 110 ± 7.4 days old). Optical stimulation was achieved by the application of blue light (473 nm) at 20 Hz with 2 ms pulses (4% duty cycle). Behavioral testing was conducted under both high laser power (4.0-4.5 mW) and low laser power (0.2-0.25 mW) conditions. Low power testing was employed to control for potential behavioral effects related to antidromic stimulation. On day 1, mice underwent OF testing and were randomly assigned to high or low power stimulation conditions. On day 2 mice were tested in the open field arena for a second time under opposite power conditions. On day 3 mice were tested in the RTPP arena for 10 minutes and were again randomly assigned to high or low power conditions. On day 4, mice were tested in the RTPP arena for 10 minutes without any stimulation to assess retention of place memory. Finally, on day 5, mice underwent RTPP under opposite power conditions (low vs high) and opposite chamber as stimulation zone (stripes vs spots) to those on Day 3.

#### Open Field Test

OF test was conducted in an opaque arena measuring 40.64 x 40.64 x 40.64 cm. Animals were placed in the center of the arena and allowed to explore freely for 10 minutes. This was followed by 5 minutes of discontinuous bilateral photostimulation at either high or low power, 5 minutes of recovery, another 5 minutes of discontinuous photostimulation at 20 Hz (at either high or low power), and a final 5 minutes of recovery. The discontinuous photostimulation consisted of three 1-minute photostimulation periods interspersed with 1-minute intervals of no photostimulation. Movement during the OF test were recorded and analyzed using ANY-maze® software (Stoelting Company, Wood Dale, IL).

#### Real-Time Place Preference

The real-time place preference (RTPP) apparatus consisted of a rectangular behavioral arena with three chambers, each measuring 24.60 x 27.94 x 27.94 cm, with a non-reflective grey background. The adjacent chambers had distinct visual cues: one chamber featured stripes and the other spots. These striped and spotted zones were randomly assigned as stimulation zones between animals. The center chamber and one of the adjacent chambers served as neutral zones, while the other adjacent chamber served as a stimulation zone. Animals received unilateral optical stimulation when in the assigned stimulation zone. Laser-on side was randomly chosen and counter-balanced between mice. Mice were also randomly assigned to have low laser power RTPP first or high laser power RTPP first. In each case, mice were given an unstimulated 10-minute trial on the day between the first and second RTPP experiment to ‘unlearn’ which side was stimulated and the second RTPP experiment stimulated the opposite chamber compared to the first RTPP experiment. For example, one mouse would have high power stimulation on the striped side on day 1, no stimulation on day 2, and low power stimulation on the spotted side on day 3. To mitigate locomotor effects with bilateral stimulation, unilateral optical stimulation was used in RTPP. Individual subjects were placed in the RTPP apparatus and allowed 10 minutes to explore both compartments. The time spent in each compartment was recorded in real-time using ANY-maze® software (Stoelting Company, Wood Dale, IL).

#### Immunohistochemistry and confocal imaging

After electrophysiological experiments, brain slices were fixed overnight in a 4% w/v paraformaldehyde (PFA) solution in phosphate buffer (PB) solution, pH 7.6 at 4°C. The fixed brain slices were then stored in phosphate buffer (PB) solution until immunostaining.

Brain slices were also collected at the conclusion of behavioral experiments. Mice were deeply anesthetized and perfused with phosphate buffer (PB), followed by a fixative solution containing 4% w/v paraformaldehyde (PFA) in PB, pH 7.6, at 4°C. Whole brains were extracted and fixed overnight in the same PFA solution. After fixation, brains were stored in PB solution until further processing. For cubic processing, 200 µm-thick sagittal slices were obtained from post-behavioral whole mouse brains using the PELCO easiSlicer™ Vibratory Tissue Slicer.

A CUBIC tissue clearing protocol ^43^ was combined with immunofluorescence staining as in^79^ for all fixed brain slices from both electrophysiological and behavioral experiments. All steps are performed at room temperature on a shaker. Slices were placed in CUBIC reagent 1 for 1-2 days; then washed in PB 3x for 1 hour each; placed in blocking solution (0.5% fish gelatin in PB) for 3 hours; placed in primary antibodies for 2-3 days; washed in PB 3x for 2 hours each; placed in secondary antibodies for 2-3 days; washed in PB 3x for 2 hours each; and placed in CUBIC reagent 2 for 2 hours before mounting onto slides (Fisherbrand 12-550-403) in reagent 2 and sealed with frame-seal incubation chambers (Thermo Scientific AB-0577) and a coverslip (Corning 2845-18).

Neurobiotin-filled patched neurons were stained with streptavidin antibody (Cy5, Invitrogen Cat#SA1011, 1:1000 or DyLight™ 405, Invitrogen Cat#21831, 1:1000). Goat anti-ChAT primary antibody (Sigma Cat#AB144P, 1:200) and donkey anti-goat Alexa Fluor™ 647 secondary antibody (Invitrogen Cat#A-21447, 1:333) was used to identify the borders of the PPN. On an example slice to show the SNr injection site, sheep anti-TH primary antibody (Novus Biologicals Cat#NB300-110, 1:1000) and donkey anti-sheep (1:100) was used to delineate the GABAergic SNr from the dopamine-rich TH+ SNc neurons. To identify the approximate location of patched cells for cell mapping, slices were imaged as tiled z-stacks using a Leica SP8AOBS++ at the Microscopy and Imaging Shared Resource core facility at Georgetown University. Additionally, to verify the injection site and optical implant placement in the brain slices from mice that underwent behavioral testing, tiled z-stacks were obtained using either the aforementioned Leica SP8AOBS++ or a Zeiss Axio Imager Z2 at the Georgetown Department of Neuroscience core imaging facility.

#### Approximate Cell Location

An image of the PPN with the pipette tip still in position was taken at the end of each cell recording and confocal images of the slices after CUBIC clearing were obtained as described above. Confocal images were overlapped with the pipette images using the streptavidin-stained patched cells (which aligned with the pipette tip images) and autofluorescence of the superior cerebellar peduncle (scp) white fibers as landmarks. The perimeter of the PPN was determined by the presence of ChAT+ neurons and generally extended from the edge of the SNr to approximately 2/3 up the scp fibers. ChAT+ neurons were identified by tdTomato expression in the ChAT-Cre mouse line and anti-ChAT staining in the Vgat-Cre and Vglut2-Cre mouse lines. Since there is variability introduced during the slicing process, we matched slices to three sagittal templates representing 1.00, 1.25, and 1.50 mm lateral to the midline in the Paxinos and Franklin’s Mouse Brain Atlas 4th edition. The most medial slice (1.00 mm lateral to the midline) characteristically had few to no fibers of the superior cerebellar peduncle at the edge of the slice where the cerebellum would tether. All neurons on this slice were considered ‘caudal’ following the typical shape of the PPN. Slices matching 1.25 and 1.50 mm from the midline had a region of densely clustered cholinergic neurons and a region of loosely dispersed cholinergic neurons. An oval ROI was created following the length and orientation of ChAT+ neuron spread. The midline of the oval demarcated the rostral and caudal regions. Overlapping the midlines allowed us to evaluate rostral-caudal differences across mice with a small margin of error, particularly along the midline. For this reason, we also mapped the location of patched cells which shows the distance from the midline and can reveal medial-lateral differences. The XY coordinates were determined using Fiji software and plotted in Igor. The color scale applied to the points represented the amplitude of the first oIPSC and ranged from 0 to 250pA so that amplitudes greater than 250pA were shown in the same maximal bright color.

#### Approximate Implant Location

To verify the location of the optical implant, anatomical landmarks for locating the PPN were identified by immunofluorescent labeled PPN cholinergic neurons. Sections containing visible optical fiber scarring were stained with anti-ChAT immunofluorescent labeling. These histological images were then visually matched to the corresponding sagittal planes of the stereotaxic atlas, based on the identified anatomical landmarks. The approximate position of the optical cannula was localized in the histological images and was in the middle-to-rostral part of the PPN.

### QUANTIFICATION AND STATISTICAL ANALYSIS

Electrophysiological traces were processed in Igor Pro 9 (Wavemetrics) and all statistical analyses were performed in GraphPad Prism 9.5. All parametric data in text is reported as mean ± standard error of the mean (SEM). Two-tailed unpaired t-tests were used to compare between two groups and one-way ANOVA followed by Dunnett’s post hoc test was used to compare three or more groups. If the equality of variances was violated (i.e. the ratio of the largest and smallest standard deviations was greater than 2 and Bartlett test was statistically significant), Welch’s ANOVA test was alternatively used. Two-way ANOVA followed by Fisher’s LSD post hoc test was performed for percent of pre-optical stimulation frequency as each comparison group was predetermined. Non-parametric data in text is reported as median [interquartile range (IQR)]. Boxplots show medians, 25th and 75th percentile as first and third quartile box edges, and 9th and 91st percentiles as whiskers. We used Mann-Whitney rank-sum test for comparing two groups and Kruskal-Wallis for multiple comparisons. As each comparison group within the Kruskal-Wallis tests was planned and stand alone, uncorrected Dunn’s post hoc test was performed to determine significance between groups. To determine if caudal GPe-connectivity is significantly greater than rostral GPe-connectivity, as expected from axon projection patterns, a one-tailed Fisher’s exact test and odds ratio effect size was performed. For correlation matrix analyses, non-parametric Spearman r was computed. Statistical details of experiments can be found in the text, figure legends, and Supplemental Table 1. Biological replicates are individual cells (n) from 6 separate mice (N=3 males and 3 females) in the electrophysiological experiments and are individual mice of both sexes in the behavioral experiments (numbers described in results text). Same sex littermates were randomly allocated to control or experimental groups and experimenters were not blinded during experiments. Statistical significance was evaluated as p < 0.05.

### Supplemental information

Document S1. Figure S1, related to Figures 2, 3, 4, & 6; Figure S2, S3, S4 related to Figure 7. Document S2. Table S1, related to Figures 2, 3, 4, 5, & 6.

Video S1. Example video of stimulating SNr axons over the PPN in an open field, related to Figure 7

Video S2. Example video of stimulating GPe axons over the PPN in an open field, related to Figure 7

Video S3. Example video of stimulating SNr axons over the PPN in the striped zone, related to Figure 7

Video S4. Example video of stimulating GPe axons over the PPN in the striped zone, related to Figure 7

---

**Figure S1.**
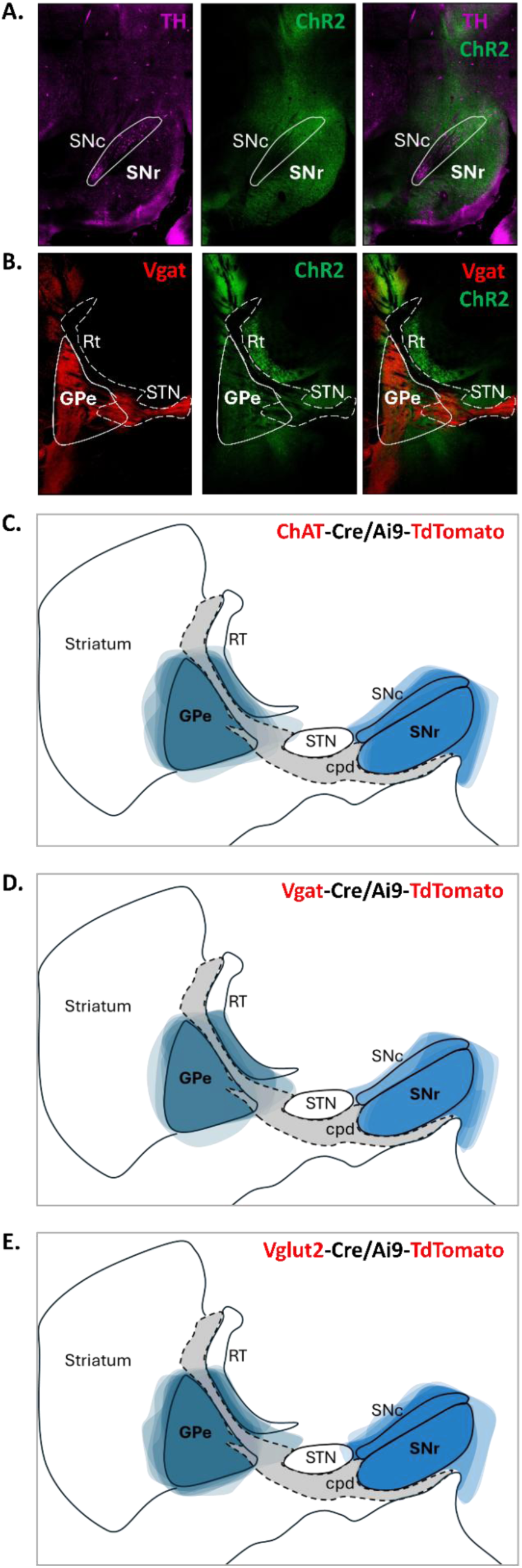
(A) Representative image of SNr injected with ChR2-EYFP. Cell bodies and axons filled with ChR2 are represented in green. Post hoc staining of tyrosine hydroxylase (TH) to label dopamine neurons in the substantia nigra *pars compacta*. TH+ neurons are represented in magenta. **(B)** Representative image of GPe injected with ChR2-EYFP in a Vgat-Cre/Ai9-TdTomato mouse. Cell bodies and axons filled with ChR2 are represented in green. Vgat+ neurons and their axonal projections are represented in red. **(C-E)** Graphical depiction of virus spread in N=6 SNr and N=6 GPe injected mice in each mouse line: **(C)** ChAT-Cre/Ai9-TdTomato, **(D)** Vgat-Cre/Ai9-TdTomato, and **(E)** Vglut2-Cre/Ai9-TdTomato mice.

**Figure S2.**
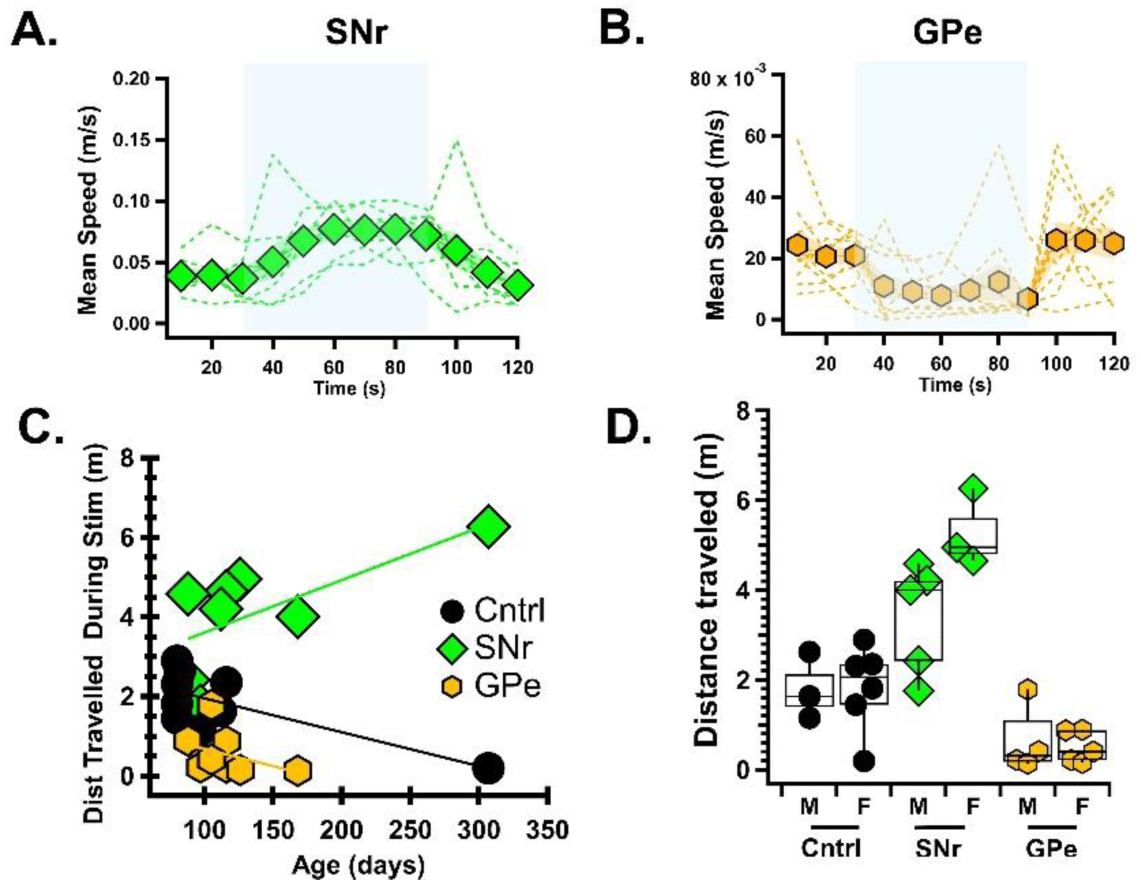
**Speed measurements across age and sex**. (A) Speed of SNr-ChR2 mice during optical stimulation in the open field. Individual mouse traces are shown as green broken lines, with the group average represented by a solid green line with diamond markers. SNr-ChR2 mice exhibited increased speed during stimulation. (B) Speed of GPe-ChR2 mice during six 1-minute optical stimulations in the open field. Individual mouse traces are shown as orange broken lines, with the group average represented by a solid orange line with hexagon markers. GPe-ChR2 mice exhibited a reduction in speed during stimulation. Data are presented as mean ± SEM.(C) Distance traveled in the open field as a function of age in SNr-ChR2 and control mice. Older SNr-ChR2 mice tend to travel further, whereas older control mice travel less. Individual mouse traces are shown. (D) Comparison of locomotion between male and female SNr-ChR2 mice in the open field, showing a trend toward increased movement in female mice with SNr stimulation.

**Figure S3.**
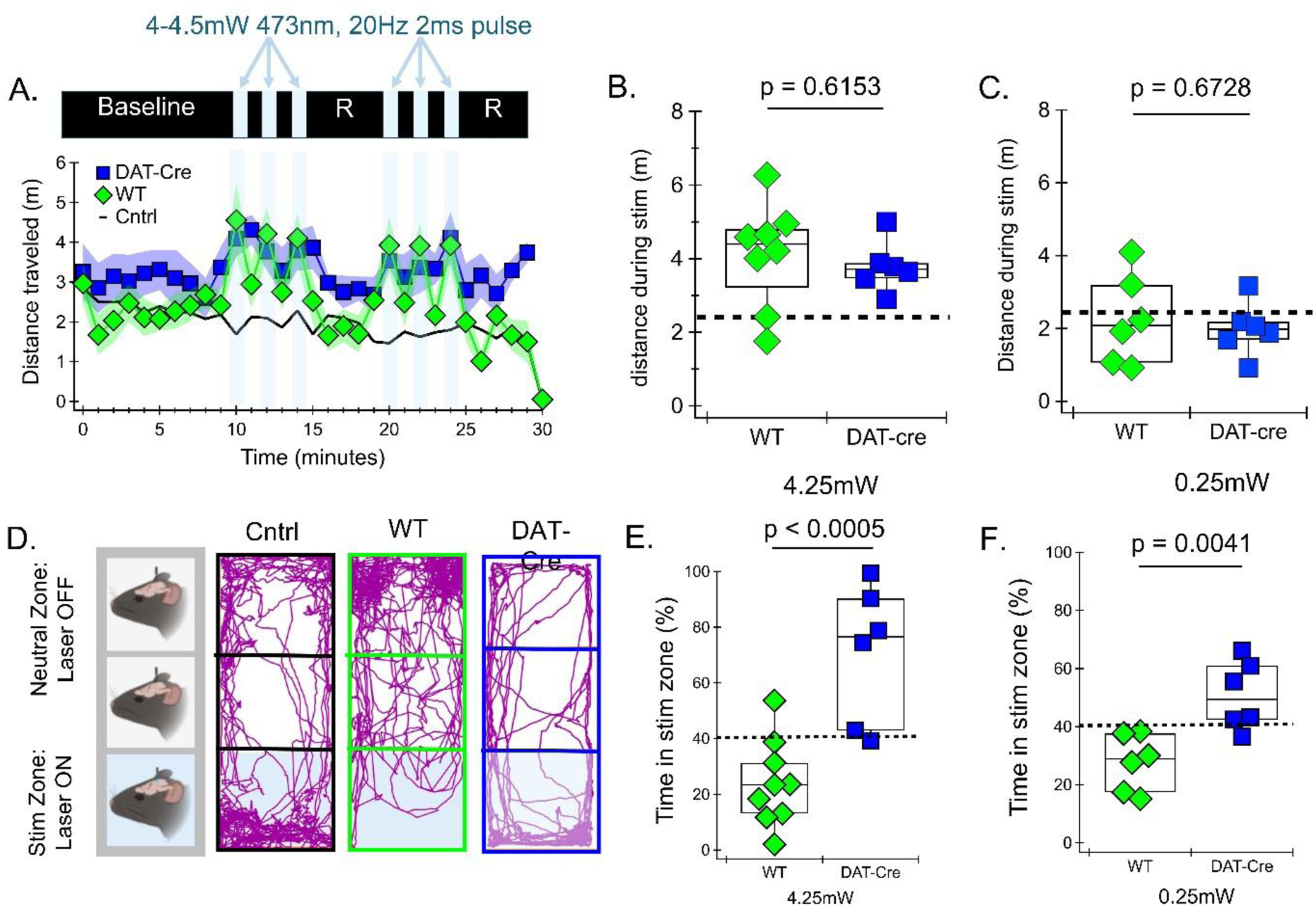
**Opposite place preference despite similar motor responses to nigral axon stimulation in the PPN of DAT-Cre and wildtype mice.** (B) Distance traveled over time in an open field during 1-minute optical stimulations (20 Hz, 4.25 mW) over the pedunculopontine nucleus (PPN). Data are shown for **N = 6** DAT-Cre mice and wildtype mice injected with ChR2 in the substantia nigra. SNr data from Figure 7. Data are presented as mean ± SEM. (C) Average distance traveled for each mouse during stimulation across six 1-minute optical stimulations under high laser power (4.25 mW). (C) Average distance traveled for each mouse during stimulation across six 1-minute optical stimulations under low laser power (0.25 mW). (D) Representative movement traces from individual mice during a real-time place preference task in a three-chamber box. Mice received continuous optical stimulation (20 Hz) of EGFP-or ChR2-expressing axons over the PPN when in the stimulation zone. (E) Percentage of time spent in the stimulation zone under high laser power (4.25 mW) for **N = 9** wildtype and **N = 6** DAT-Cre mice injected with ChR2 in the substantia nigra. (F) Percentage of time spent in the stimulation zone under low laser power (0.25 mW) for the same groups.

**Figure S4.**
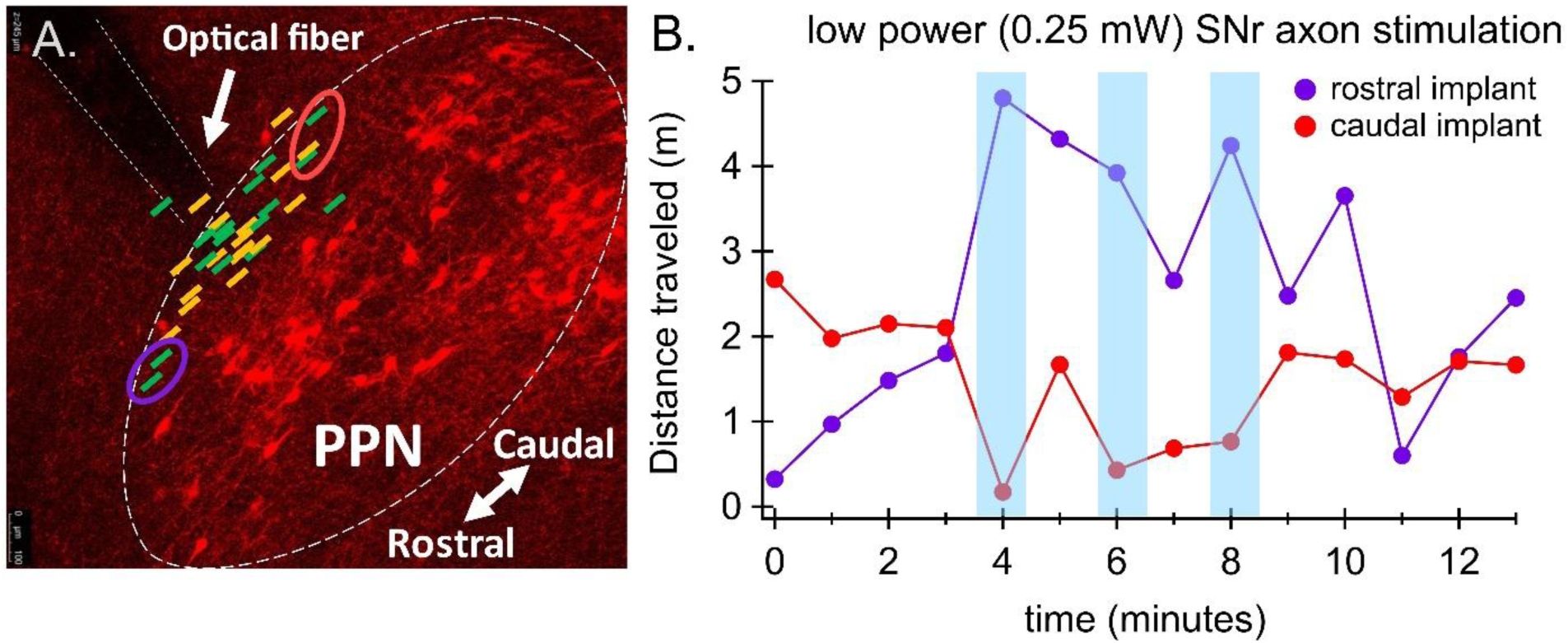
Rostral vs caudal implant site alters SNr axon stimulation effects. (A) Implant location (same as Figure 7B) with most rostral (purple) and most caudal (red) bilateral implant pairs circled for SNr axon stimulaton. (B) Distance traveled in open field in mouse with rostral implants (purple) and mouse with caudal implants (red).

**Table 1.** Statistical details of exact p-values and critical values for SNr input to ChAT+ PPN neurons.

| Parameter | Statistical Test | p-value | ChAT+ |  |
| --- | --- | --- | --- | --- |
|  |  |  | Rostral | Caudal |
| **oIPSC Amplitude** | unpaired t-test, $t=0.9041$, $df=33$ | 0.3725 | $n=15$, $92.6 \pm 18$ pA | $n=20$, $115.2 \pm 17$ pA |
| **PPR** | unpaired t-test, $t=0.1082$, $df=33$ | 0.9145 | $n=15$, $0.96 \pm 0.04$ | $n=20$, $0.97 \pm 0.05$ |
| **Avg membrane potential** | Mann Whitney, $U=87$ | 0.4706 | $n=11$, $-54.24$ ($-56.94$ to $-50.20$) mV | $n=19$, $-53.00$ ($-55.38$ to $-48.84$) mV |
| **% Pre-Opto Frq** | Repeated Measures 2way ANOVA<br>Stimulation: $F(1, 35)=81.9$<br>Region: $F(1, 35)=0.002052$<br>Interaction: $F(1, 35)=3.273$ | $<0.0001$<br>0.9641<br>0.0790 | $n=14$<br>During: $35.62 \pm 8.869$ %<br>Post: $136.1 \pm 7.803$ % | $n=23$<br>During: $52.11 \pm 8.432$ %<br>Post: $119.1 \pm 4.614$ % |
| **ΔFrq During Opto** | Mann Whitney, $U=131$ | 0.3602 | $n=14$, $-2.47$ ($-3.37$ to $-1.44$) Hz | $n=23$, $-2.10$ ($-3.11$ to $-0.84$) Hz |
| **ΔRebound Frq** | Mann Whitney, $U=79$ | 0.0142 | $n=14$, $1.64$ ($1.00$ to $2.17$) Hz | $n=22$, $0.83$ ($0.42$ to $1.14$) Hz |

**Table 2.** Statistical details of exact p-values and critical values for SNr input to Vgat+ PPN neurons.

| Parameter | Statistical Test | p-value | Vgat+ |  |
| --- | --- | --- | --- | --- |
|  |  |  | Rostral | Caudal |
| **oIPSC Amplitude** | Mann Whitney, $U=50$ | 0.6005 | $n=9$, $76.5$ ($27.0$ to $209.9$) pA | $n=13$, $59.7$ ($37.7$ to $269.7$) pA |
| **PPR** | Mann Whitney, $U=48$ | 0.5123 | $n=9$, $1.01$ ($0.88$ to $1.16$) | $n=13$, $1.06$ ($0.96$ to $1.15$); |
| **Avg membrane potential** | unpaired t-test, $t=2.468$, $df=13$ | 0.0282 | $n=5$, $-57.91 \pm 1.630$ mV | $n=10$, $-52.00 \pm 1.472$ mV |
| **% Pre-Opto Frq** | Repeated Measures 2way ANOVA<br>Stimulation: $F(1, 24)=44.63$<br>Region: $F(1, 24)=0.09642$<br>Interaction: $F(1, 24)=0.8841$ | $<0.0001$<br>0.7588<br>0.3565 | $n=7$<br>During: $37.91 \pm 15.91$ %<br>Post: $100.0 \pm 14.57$ % | $n=19$<br>During: $35.62 \pm 8.869$ %<br>Post: $113.2 \pm 5.187$ % |
| **ΔFrq During Opto** | Mann Whitney, $U=55$ | 0.5336 | $n=7$, $-5.03$ ($-5.55$ to $-1.34$) Hz | $n=19$, $-4.29$ ($-11.02$ to $-2.94$) Hz |
| **ΔRebound Frq** | Mann Whitney, $U=63$ | $>0.9999$ | $n=7$, $0.80$ ($-1.31$ to $2.33$) Hz | $n=18$, $0.59$ ($-2.35$ to $1.47$) Hz |

**Table 3.** Statistical details of exact p-values and critical values for SNr input to Vglut2+ PPN neurons.

| Parameter | Statistical Test | p-value | Vglut2+ |  |
| --- | --- | --- | --- | --- |
|  |  |  | Rostral | Caudal |
| **oIPSC Amplitude** | Mann Whitney, U=29 | 0.0035 | n=13, 61.6 (19.2 to 113.9) pA | n=13, 313.5 (143.8 to 729.1) pA |
| **PPR** | Mann Whitney, U=77 | 0.7241 | n=13, 1.07 (0.86 to 1.38) | n=13, 0.92 (0.89 to 1.02) |
| **Avg membrane potential** | unpaired t-test, t=0.3838, df=25 | 0.7043 | n=11, -53.03 ± 1.815 | n=16, -52.09 ± 1.601 |
| **% Pre-Opto Frq** | Repeated Measures 2way ANOVA<br>Stimulation: F(1,28)=133.6<br>Region: F(1, 28)=0.5214<br>Interaction: F(1,28)=0.1765 | <0.0001<br>0.4762<br>0.6776 | n=13<br>During: 32.62 ± 10.21 %<br>Post: 100.3 ± 6.113 % | n=17<br>During: 24.71 ± 7.297 %<br>Post: 97.48 ± 2.228 % |
| **ΔFrq During Opto** | Mann Whitney, U=55 | 0.0343 | n=13, -3.21 (-12.21 to -1.95) Hz | n=17, -14.08 (-20.04 to -6.07) Hz |
| **ΔRebound Frq** | Mann Whitney, U=88 | 0.3627 | n=13, 0.37 (-0.027 to 0.48) Hz | n=17, -0.23 (-0.82 to 1.08) Hz |

**Table 4.** Statistical details of exact p-values and critical values for GPe input to ChAT+, Vgat+, and Vglut2+ PPN neurons.

| Parameter | Statistical Test | p-value | ChAT+ | Vgat+ | Vglut2+ |
| --- | --- | --- | --- | --- | --- |
| **oIPSC Amplitude** | Kruskal-Wallis, H(2,37)=3.050 | 0.2176 | n=6, 22.6 (9.75 to 50.20) pA | n=19, 24.77 (6.72 to 54.17) pA | n=15, 65.71 (14.50 to 133.4) pA |
| **PPR** | Mann Whitney, U=76 | 0.0206 |  | n=19, 0.97 (0.76 to 1.2) | n=15, 0.81 (0.55 to 0.93) |
| **Avg membrane potential** | Kruskal-Wallis, H(2,69)=2.963 | 0.2273 | n=25, -54.16 (-58.05 to -51.10) mV | n=18, -54.28 (-55.47 to -48.77) mV | n=29, -55.32 (-57.97 to -52.28) mV |
| **% Pre-Opto Frq** | Repeated Measures 2way ANOVA<br>Stimulation: F(1, 69)=25.46<br>Cell type: F(1, 69)=4.444<br>Interaction: F(1, 69)=2.805 | <0.0001<br>0.0153<br>0.0674 | n=29<br>During: 92.39 ± 7.59 %<br>Post: 104.10 ± 5.23 % | n=18<br>During: 57.70 ± 10.11 %<br>Post: 91.81 ± 6.98 % | n=24<br>During: 64.15 ± 7.63 %<br>Post: 104.7 ± 5.39 % |
|  | Uncorrected Fisher's LSD, df=138<br>t=2.971<br>t=3.223<br>t=0.6177<br>t=0.1407<br>t=1.076<br>t=1.236 | 0.0035<br>0.0016<br>0.5378<br>0.8883<br>0.2838<br>0.2184 | During:<br>ChAT+ vs. Vglut2+: 28.24 (CI 9.446 to 47.03)<br>ChAT+ vs. Vgat+: 34.69 (CI 13.41 to 55.97)<br>Vglut2+ vs. Vgat+: 6.454 (CI -14.21 to 27.11)<br>Post:<br>ChAT+ vs. Vglut2+: -1.337 (CI -20.13 to 17.45)<br>ChAT+ vs. Vgat+: 11.58 (CI -9.702 to 32.86)<br>Vglut2+ vs. Vgat+: 12.92 (CI -7.741 to 33.58) |  |  |
| **ΔFrq During Opto** | Kruskal-Wallis, H(2,69)=17.21 | 0.0002 | n=25, -0.16 (-0.90 to 0.03) Hz | n=18, -1.92 (-3.85 to -0.60) Hz | n=29, -0.89 (-3.28 to -0.43) Hz |
|  | Uncorrected Dunn's test | 0.0009<br>0.0002<br>0.4020 | ChAT+ vs. Vglut2+: Z=3.332<br>ChAT+ vs. Vgat+: Z=3.755<br>Vglut2+ vs. Vgat+: Z=0.8380 |  |  |
| **ΔRebound Frq** | Kruskal-Wallis, H(2,69)=17.21 | 0.6772 | n=8, 0.1041 (-0.52 to 0.41) Hz | n=19, 0.07082 (-0.36 to 0.57) Hz | n=7, -0.1421 (-2.58 to 0.71) Hz |

